# Evolutionary Recoding of Olfactory Sensory Neurons

**DOI:** 10.64898/2026.06.12.731899

**Authors:** Gwénaëlle Bontonou, Tess Baticle, Sarah Hume, Tane Kafle, Batool Mahmoud, Vaia Vlachou, Matthew Day, J. Roman Arguello

**Affiliations:** Department of Ecology & Evolution, Faculty of Biology and Medicine, University of Lausanne, Lausanne, Switzerland; School of Biological and Behavioural Sciences, Queen Mary University of London, London, UK; Centre for Evolutionary and Functional Genomics, Queen Mary University of London, London, UK

## Abstract

Odorant receptors (*Or*s) are the interface between an animal’s nervous system and its olfactory environment. In insect genomes, the *Or*s are often the largest gene family, and their rapid duplications and deletions result in extensive copy-number differences between species. Because olfactory sensory neurons (OSNs) typically express only one *Or* (the so-called ‘one-receptor one-neuron rule’), this dynamism at the level of *Or* genes raises fundamental questions regarding their cellular regulation: How do new *Or* duplicates gain their own neuron-specific expression? Such Or-OSN changes are thought to provide a key evolutionary path for modifying olfactory perception and related behaviours, but the absence of examples of these transitions has prevented an understanding of how they occur. Using a highly duplicated *Drosophila Or* subfamily (the Or67a subfamily) as a model system, we discovered parallel instances of *Or67a* duplicates gaining new OSN expression and reconstructed their evolutionary histories. Functional work in *D. suzukii*, a species with two novel *Or67a*-expressing OSN populations, revealed that their *Or67a* expression has arisen in preexisting OSNs, which have lost their ancestral *Or*s. As a result, these neurons were recoded and acquired new olfactory identities, thereby demonstrating the diversification of an OSN repertoire without the invention of developmentally new OSN lineages.

## Introduction

Olfaction and olfactory-guided behaviours are incredibly variable. A key driver of these differences is the evolution of new olfactory sensory neuron (OSN) functions. Understanding how these new functions emerge is of fundamental importance and, in the cases of managing pests and disease vectors, of applied importance too^1–4^.

Analogous to vertebrates, insect OSNs predominantly express a single type of Odorant receptor (*Or*) gene^5^. Though reports of *Or* co-expression are rising, their pattern of expression has been referred to as “one receptor-one neuron”^6–8^. The volatile molecules that insect Or proteins bind define the physiological tuning of each OSN population, and their ligand-gated activation initiates behavioural responses that are ecologically important and often species-specific^2,9–11^. The gene families that encode Ors are among the largest in insect genomes and their members frequently evolve rapidly^2,12–15^. In addition to low protein similarity among orthologs and frequently unidentifiable homologs across insect taxa, their fast evolution is fostered by high rates of copy number changes. These mutations result in significant differences in the Or family sizes and functions across species^7,16–19^.

That each OSN population typically expresses only a single *Or* gene is striking considering how frequently they duplicate, and raises fundamental questions over how new *Or* copies become expressed singularly in their own OSN population. Peripheral olfactory nervous systems are presumably diversified through these ongoing processes. How this occurs – and how often – remains unknown. Addressing these questions requires reconstructing the evolutionary histories of *Or* duplicates at the genomic, OSN, and physiological levels. To date, however, most evolutionary studies have focused primarily on quantifying *Or* copy number changes between species, leaving transitions in their OSN expression and their impacts on olfactory perception largely unexplored.

We used the most duplicated *Or* subfamily in the *D. melanogaster* species group, the *Or67a* subfamily, to investigate these questions. We reasoned that its elevated rate of duplication would maximise our chances of capturing hitherto undocumented transitions in expression and function for its recent duplicates. Comparative analyses across 18 *Drosophila* species revealed that, contrary to the predominant “one receptor-one neuron rule”, most *Or67a* duplicates are co-expressed with one or more *Or67a* paralogs in a homologous OSN population. Despite broad co-expression, we discovered two *Or67a* duplicates that have escaped their ancestral expression and independently gained distinct OSN expression. Focusing on *D. suzukii*, a species with both new *Or67a*-expressing OSNs, we demonstrate that the recipient OSNs were preexisting neuron populations. Strikingly, in both cases, the gain of *Or67a* expression has corresponded with the loss of the neurons’ ancestrally expressed *Or*s, revealing parallel receptor replacements. Using extracellular recordings, we show that *D. suzukii*’s *Or67a* copies have functionally diversified and that the *Or* replacements have resulted in a “recoding” of the two recipient OSN populations. Together, our data provide the first evolutionary reconstruction of how duplicated *Or*s diversify olfactory perception through the recoding of existing OSNs.

## Results

### The *Or67a* subfamily is mobile

The size of the *Or67a* subfamily is known to vary across members of the *D. melanogaster* species group, but the evolutionary history underlying the diversity in copy number has not been investigated^7,19–21^. To reconstruct the subfamily’s evolutionary past, we first inspected the homologous chromosomal regions that contain an *Or67a* gene (or gene fragment) in 15 species from the *D. melangaster* species group and three outgroup species (Figure 1A, Figures S1-10, Files S1-2).

**Figure 1:**
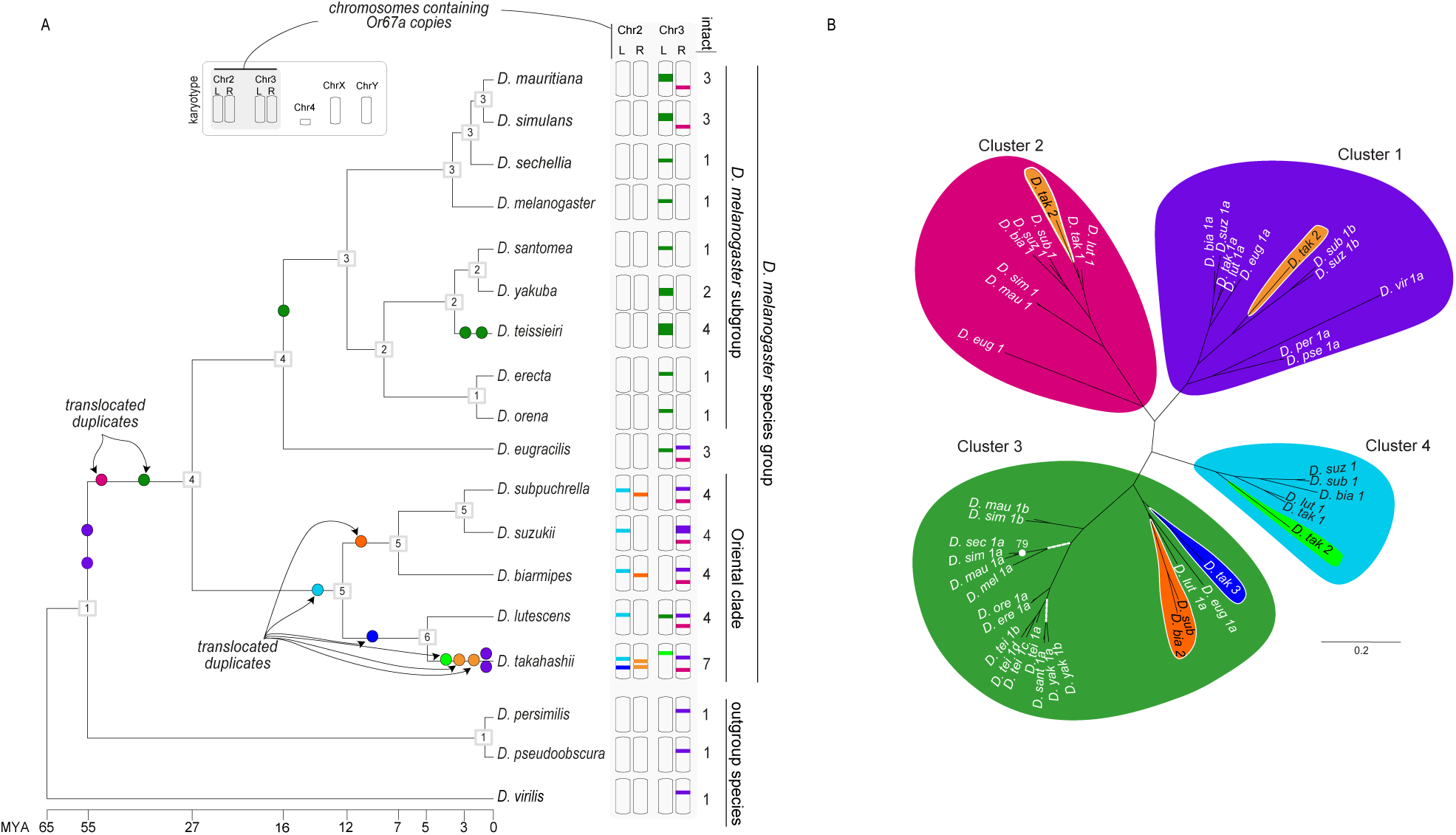
Evolution of the *Or67a* subfamily within the *Drosophila melanogaster* species group. **A)** Number and chromosomal locations of intact *Or67a* paralogs. For simplicity, only gene duplications are mapped onto the species tree (not to scale), with translocated duplications annotated (see Figure S1 for *Or67a* gains and losses). The number of genes inferred at ancestral nodes is indicated in boxes. Chromosomal locations of *Or67a* paralogs are shown at the tree tips, along with the number of intact copies. **B)** Protein tree of the Or67a subfamily inferred from 46 intact sequences across 18 *Drosophila* species. All nodes have posterior support >80, unless noted by a white dot and the node’s support. White hashed branches indicate significantly high rates of protein evolution were detected (elevated dN/dS). Orthologous genes are shaded in different colours and correspond to the colours in panel A: purple/pink (chromosome 3R), green (3L), yellow/orange (2R), and dark/light blue (2L). Species names are abbreviated using the first four letters of the genus and species. Scale bar: 0.2 amino acid substitutions per site.

Our reannotations and microsynteny analyses identified 46 functional *Or67a* copies and 9 pseudogenes across the 18 *Drosophila* species (Figure 1A, Figure S1). When we placed these analyses within a phylogenetic and chromosomal context, we discovered that *Or67a* was ancestrally a single-copy gene on chromosome 3R (corresponding to Muller element E in *D. virilis*, *D. persimilis* and *D. pseudoobscura*) but has since been mobilised by recurrent translocated duplication events, eight in total (Figure 1A). Most of these events were interchromosomal and have resulted in the seeding of both arms of chromosomes 2 and 3 with new *Or67a* copies.

We inferred that the mobilisation of *Or67a* copies began with two translocated duplication events in the common ancestor of the *D. melanogaster* subgroup and the Oriental Clade, which seeded a more distal region of chromosomes 3R and 3L. These two translocated copies have only partially been retained in the extant species, with the genomes from the *D. melanogaster* subgroup predominantly retaining the chromosome 3L copy and the Oriental Clade retaining the chromosome 3R copy. No further translocated duplicates were found in the *D. melanogaster* subgroup, but the frequency of these events increased within the Oriental Clade, particularly along the branch leading to *D. takahashii*. Six additional translocated duplicates further seeded chromosome 3L and both arms of chromosome 2 (Figure 1A).

The genomic intervals containing *Or67a* paralogs harbour multiple transposable elements (TEs) and TE remnants, as well as indels, and two inversions, suggesting these regions have high mutation rates and may be unstable (Figures S2-10). TEs have been associated with translocated duplications due to the ectopic recombination they can foster, and through their capture of flanking DNA when active^22,22–26^. Despite their prevalence, we did not observe obvious signatures of ectopic recombination, however: The families that the TE fragments belong to are variable across the *Or67a*-containing loci, and when investigating the *Or67a*-containing regions of *D. suzukii* and *D. takahashii* - the two species that qualitatively appeared to have the most TEs nearby *Or67a* genes - we found that their level of association was not significantly different compared to genomic background expectations (Methods). We also did not identify any active TEs that have been described to capture flanking DNA (e.g. Helitrons^23,27^). As a result, the causal agent(s) responsible for the mobilisation of *Or67a* copies remain unclear.

The dynamic turnover of *Or67a* genes raised questions about the roles of selective constraint and adaptive selection throughout the subfamily’s history. To investigate the types of evolutionary forces acting on the subfamily, we first inferred a protein tree using the intact *Or67a* copies (Figure 1B). This well-supported protein tree reflects its history of recurrent translocations, with the four prominent clusters each containing orthologs of the oldest translocation events (and, in some cases, additional tandem duplicates). Nested within each cluster are the younger duplicated translocations. Hereafter, we use these clusters to name the *Or67a* members (e.g., *Or67a.C1* or *Or67a.C2*; for tandem duplicates within a cluster, we additionally letter it: *Or67a.C1.1<u>a</u>* and *Or67a C1.1<u>b</u>*). Using this Or67a protein tree, we tested evolutionary models of protein evolution by fitting rates of amino-acid-changing (dN) and silent (dS) substitutions along its branches. Among the models we investigated, the best-fitting models showed strong selective constraint across nearly all branches (dN/dS < 0.30), providing evidence that the intact copies are functional receptors (File S3). The two branches that were exceptions had dN/dS values greater than one, consistent with adaptive protein changes. One of these branches is ancestral to the *Or67a.C3.1a* ortholog that is shared by *D. maurtiana, D. simulans, D. sechellia,* and *D. melanogaster*, and is consistent with our previous findings^7^ (dN/dS = 1.2). The second branch is ancestral to the *Or67a.C3.1a* ortholog shared by *D. santomea* and *D. yakuba* (and *D. yakuba*’s *Or67a.C3.1b*; dN/dS = 2.9). These findings suggest that Cluster 3 is a repeated target of adaptive evolution within the *D. melanogaster* subgroup.

### Typically, *Or67a* duplicates are co-expressed, but two have gained new OSN expression

The evolutionary steps that result in a newly duplicated *Or* acquiring the “one receptor-one neuron” pattern of OSN expression remain largely unknown because transitional examples haven’t been identified. The numerous duplication events that have occurred in the *Or67a* subfamily provided an outstanding opportunity to investigate this question. We began by examining the expression of paralogs belonging to six species in the *D. melanogaster* subgroup (Figures 2A-C; Methods). These experiments confirmed the co-expression of the three *Or67a* copies in the *D. simulans* and *D. mauritiana* genomes (*Or67a.C1.1a, Or67a.C1.1b, Or67a.C2*), which had been visualised using standard FISH^7^. Using an additional probe that was designed for *D. sechellia*’s pseudogenised *Or67a.C2* copy, we found that its expression co-localised in the same OSN as its single intact copy, *Or67a.C3.1a* (Figure 2C). This result provides additional support that the three *Or67a* copies that are conserved in *D. simulans’* and *D. mauritiana’s* genomes were co-expressed ancestrally in a neuron population named ab10A, and that *D. melanogaster* and *D. sechellia* have independently lost two of them (*Or67a.C1.1b* and *Or67a.C2*)^7^.

**Figure. 2:**
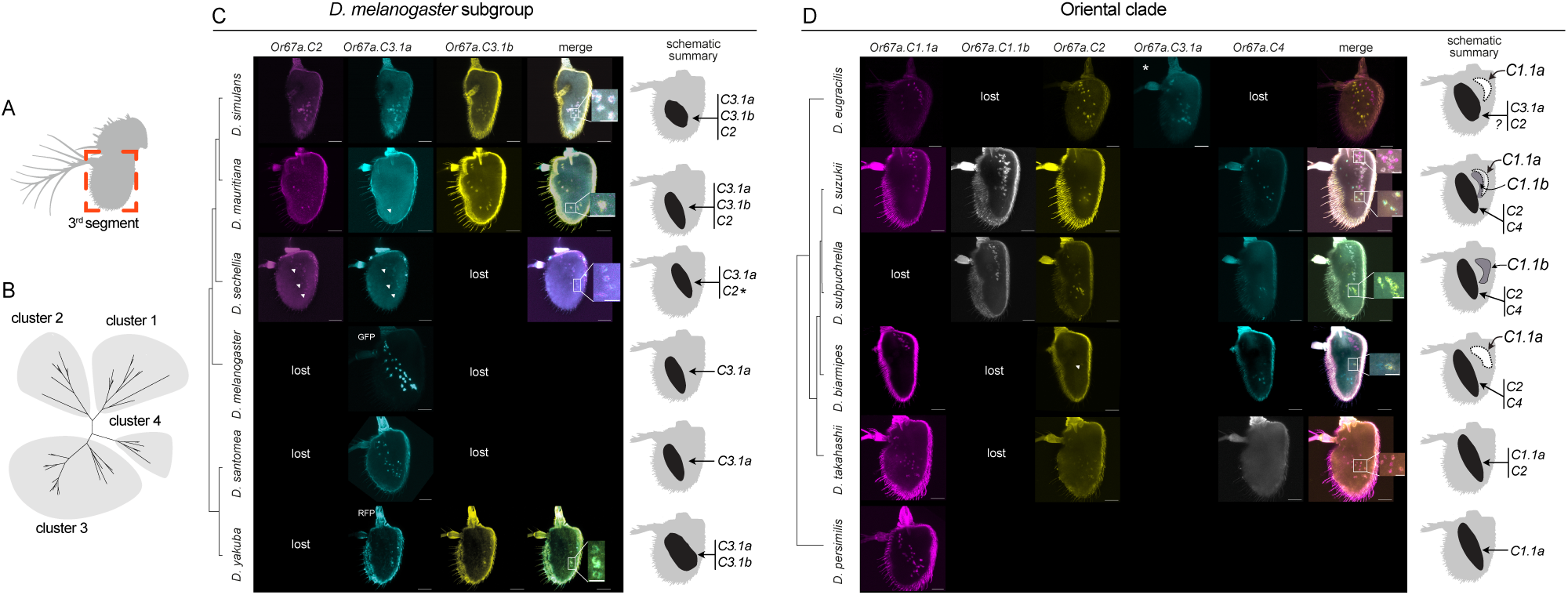
Antennal expression of *Or67a* genes. **A)** *Drosophila* antenna schematic highlighting the third segment, the focus of experiments in panels C-E. **B)** The Or67a protein tree for reference with panels C-E. **C)** Spatial patterns of *Or67a* paralog expression for species in the *D. melanogaster* subgroup. HCR-FISH was used to visualise *Or67a* expression for all species except *D. melanogaster*, for which a preexisting *Or67a.C3.1a*-GFP reporter line^30^ was imaged, and *D. yakuba*, for which an *Or67a.C3.1a*-RFP reporter line was imaged in combination with a standard *Or67a.C3.1b* FISH probe (Methods). Astrix next to *D. sechellia*’s *Or67.C2* copy indicates a pseudogene. Data are organised by the species’ relationships (tree in left margin, not to scale). **D)** HCR FISH results displaying the spatial patterns of *Or67a* paralog expression for species in the Oriental Clade (see Figure S11 for HCR FISH results that include the outgroup species *D. pseudoobscura* and *D. virilis*). The asterisk for *D. eugracilis*’s *Or67a.C3.1a* indicates single probe labelling in an antenna different from the *Or67a.C1.1a* and *Or67a.C4* paralogs (for unknown reasons, co-labelling with the three probes was not successful). Data are organised by the species’ relationships (tree in left margin, not to scale). For panels C and D, white arrowheads indicate faintly labelled cells; scale bars: Full antennae: 30 µm; enlarged view: 10 µm.

Next, we examined *Or67a* expression in a more distantly related species pair, *D. santomea* and *D. yakuba*, which diverged from a common ancestor ∼1 million years ago (MYA)^28^ and from the common ancestor of *D. melanoagaster, D. simulans* and *D. mauritiana* ∼15 MYA^29^. The ancestral *Or67a* ortholog of *D. santomea* and *D. yakub*a, *Or67a.C3.1*, carries the signature of adaptive protein changes (above) and duplicated tandemly twice in the genome of *D. yakuba* (Figure 2C; Figure S3). The *Or67a*-expressing OSNs in both species were located within a similar antennal domain to that found in *D. melanogaster* and the *D. simulans* trio, and again revealed co-expression of *D. yakuba*’s tandem duplicate copies (*Or67a.C3.1a*, *Or67a.C3.1b*; Figure 2C). Collectively, these data suggest that species from the *D. melanogaster* subgroup express their singular or multiple *Or67a* copies in a single OSN population that is homologous with *D. melanogaster*’s ab10A population.

Because the Oriental Clade has experienced elevated rates of *Or67a* duplication, we wondered if this might have fostered the evolution of more diverse expression patterns than those found in the *D. melanogaster* subgroup. We broadened our HCR-FISH experiments to include four species from the Oriental clade (*D. suzukii*, *D. subpulchrella*, *D. biarmipes*, and *D. takahashii*), *D. eugracillis* (a species that falls between the *D. melanogaster* subgroup and the Oriental Clade), and three additional outgroup species to both clades, *D. persimilis, D. pseudoobscura* and *D. virilis* (Figure 2D, Figure S11). Remarkably, though most *Or67a* copies were again found to be co-expressed in OSNs within an ab10A-like antennal domain, we found two receptors, *Or67a.C1.1a* and *Or67a.C1.1b*, have gained expression in what appeared to be distinct OSNs: *D. eugracilis*, *D. subpulchrella*, and *D. biarmipies*, each have one novel *Or67a*-expressing OSN population, while *D. suzukii* has two (Figure 2D).

### *Or67a* co-expression occurs in OSNs homologous to ab10A

We have so far assumed that the ancestral-like *Or67a*-expressing OSNs are homologous to *D. melanogaster*’s ab10A population due to their similar antennal locations. To test this assumption, we designed a co-labelling HCR-FISH experiment that was based on the complete OSN-sensilla map described in *D. melanogaster*^5,30^. Because this map describes which OSN are paired together within each class of sensilla, we knew that *D. melanogaster*’s *Or67a*-expressing OSNs are housed within sensilla named ab10. ab10 also contain an OSN population that co-expresses *Or49a* and *Or85f* (another exception to the “one receptor-one neuron” pattern; Figure 3A). If this OSN:sensilla association has remained evolutionarily conserved, we would expect to observe the pairing of the *Or67a*- and *Or49a-* and/or *Or85f-expressing* OSNs across species reflecting their containment in the same sensilla. Consistent with this expectation, we found the OSNs labelled by these probe combinations are immediately adjacent in the Oriental Clade species, in *D. eugracilis*, and in the outgroup species *D. persimilis* and *D. pseudoobscura* (Figure 3A,B; Figure S11). In contrast, the *Or67a.C1.1a-* and *Or67a.C1.1b*-expressing OSN that are outside of the ancestral domain of expression do not pair with either *Or49a-* or *Or85f*-expressing OSNs (Figure 3A,B). These results indicate that the ab10A population has not expanded into different antennal regions in *D. eugracils*, *D. suzukii*, or *D. subpulchrella* (*Or49a* and *Or85f* labelling in *D. biarmipies* was unsuccessful). Instead, the *Or67a.C1.1a* and *Or67a.C1.1b* copies in these latter species have escaped the ancestral OSN expression shared by their paralogs. Together with our previous functional studies of co-expressed *Or67a* copies^7^, these observations reveal that ab10A OSNs have been an incubator for odorant receptor diversification, occasionally spawning its duplicates into different cells.

**Figure 3:**
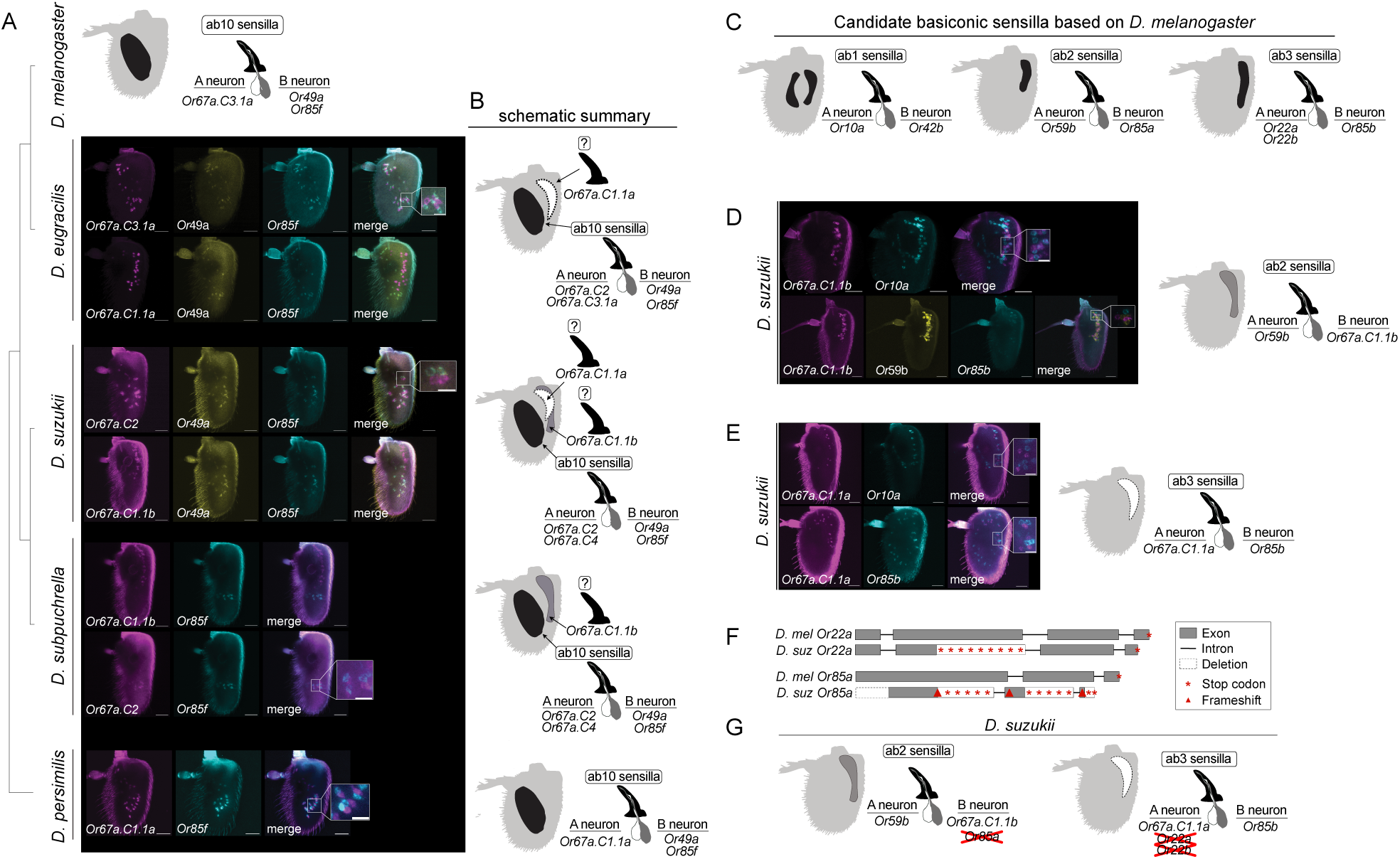
Identification of ancestral and new Or67a-expressing OSNs. **A)** Top: schematic of the D. melanogaster’s ab10 sensilla and the OSN that they contain5,28. Below: Results of co-labelling experiments using HCR-FISH probes designed to test for the pairing of Or67a expression in combination with Or85f and/or Or49a within ab10 sensilla (D. subpuchrella and D. persimilis do not have an Or49a ortholog; see Figure S11 for HCR FISH results that include the additional outgroup species D. pseudoobscura and D. virilis). **B)** Schematic summaries of the HCR-FISH results from panel A. **C)** Schematic of the D. melanogaster’s ab1-3 sensilla and the Ors that their OSN express5,28. This sensilla:Or map that was used for the test for the neuronal identity for Or67a.C1.1b expression in D. suzukii. **D)** Co-labelling experiments in D. suzukii antennae based on the sensilla:Or map in panel C, revealing Or67a.C1.1b is expressed in ab3A neurons. A schematic summarizing of the findings is shown on the right. White arrowheads indicate faintly labelled cells. **E)** Co-labelling experiments in D. suzukii antennae based on the sensilla:Or map in panel C, revealing Or67a.C1.1a is expressed in ab2A neurons. A schematic summarizing of the findings is shown on the right. **F)** Simplified illustration of D.suzukii’s Or22a and Or85a pseudogenes, which have been replaced by Or67a.C1.1a and Or67a.C1.1b, respectively. **G)** Schematic summary of the results based on panels C-F. Scale bars: Full antennae: 30 µm, Zoomed-in area: 10 µm.

### *Or67a C1.1a and C1.1b* have independently gained new expression in different OSNs

What are the identities of the neurons with the new *Or67a.C1.1a* and *Or67a.C1.1b* expression? Their cell bodies lie in a medioproximal domain of the antennae’s third segment, superficial to a cluster of sensilla named “large basiconic” based on their morphology. Because *D. melanogaster* also has this class of sensilla in approximately the same antennal domain, we speculated that *Or67a.C1.1a’s* and *Or67a.C1.1b’s* gain of expression may have occurred in the neurons that they contain. Drawing again on *D. melanogaster*’s OSN:sensilla map^5,30^, we identified three candidate sensilla types that at least partially overlap the medioproximal domain of interest: ab1, ab2, and ab3 sensilla. Each of these sensilla contain two OSNs: ab1 contains *Or10a*- and *Or42b*-expressing OSN, ab2 contains *Or59b*- and *Or85a*-expressing OSN, and ab3 contains *Or85b*- and *Or22a/22b*-coexpressing OSN (Figure 3C).

We again carried out HCR-FISH co-labelling experiments, this time to test if the *Or67a.C1.1a-* or *Or67a.C1.1b*-expressing OSN pair with (or co-localise in) OSNs labelled by probes that are diagnostic for the three sensilla types. For this work, we focused on *D. suzukii* because it has both new *Or67a*-expressing OSN populations. We began by examining the location of its *Or67a.C1.1b*-expressing OSNs relative to those labelled by *Or10a* (diagnostic for ab1 sensilla), *Or59b* (diagnostic for ab2 sensilla), and *Or85b* (diagnostic for ab3 sensilla) probes. Of the three, the OSNs expressing *Or59b* unambiguously matched the distribution of those expressing *Or67a.C1.1b*. In these experiments, the signal from the two probes did not co-localise but instead labelled OSNs that are immediately adjacent, indicating that *Or67a.C1.1b*- and *Or59b*-expressing OSNs are neighbouring neurons contained in ab2 sensilla and that *Or67a.C1.1b* is expressed in the B neuron (ab3B; Figure 3D).

With ab1 and ab3 sensilla the remaining candidates, we next carried out co-labelling experiments with the *Or67a.C1.1a* probe and either an *Or10a* (diagnostic for ab1 sensilla) or *Or85b* (diagnostic for ab3 sensilla) probe. While both *Or10a*- and *Or85b*-expressing OSN populations overlap in the medioproximal domain, they also extend more distolaterally than the *C1.1a* population (Figure 3E). We did not find any pairing (or co-expression) between *Or67a.C1.1a*- and *Or10a*-expressing neurons, but a subset of *Or85b*-expressing OSN that are in the medioproximal domain do tightly pair with the *Or67a.C1.1a*-expressing neurons. This indicates that *Or67a.C1.1a* has gained expression in the A neurons of ab3 sensilla (ab3A) and also suggests that the ab3B neurons contain at least two subpopulations, one that pairs with the *Or67a.C1.1a*-expressing OSN and another uncharacterized distolateral subpopulation (a possibility previously suspected based on physiological recordings^31^).

*Or67a.C1.1a’s* and *Or67a.C1.1b*’s gain of expression would presumably have led to the co-expression of multiple *Or*s in ab2B and ab3A neurons. However, the ancestrally expressed receptors in ab2B and ab3A neurons, *Or22a* and *Or85a*, respectively, have been pseudogenised in *D. suzukii*^20^. Manual inspection of the *Or22a* annotation, and its reannotation in seven *D. suzukii* genomes, supports it being a fixed pseudogene. We found that it has also been lost in other Oriental Clade species and *D. eugracillis* (File S4; We note that *Or22b* – a paralog of *Or22a*, with which it co-expresses in some *D. melanogaster* subgroup species – does not exist in the Oriental Clade; Figure S12). An analogous analysis of *Or85a* revealed that it too is a fixed pseudogene within *D. suzukii* but is intact in other Oriental Clade species (File S5). We therefore uncovered parallel replacements of ancestrally expressed *Or*s with new duplicated copies of the distantly related *Or67a* subfamily.

Despite *Or67a.C1.1a* and *Or67a.C1.1b* being the only two *Or67a* copies to have gained new OSN expression, the pattern of their changes escapes a parsimonious explanation. We can infer that the ab10A-like OSN was ancestral because it is shared by all species, including the outgroup species *D. persimilis* and *D. pseudoobscura* (Figure 2D, Figure S11). The distinct *Or67a.C1.1b*-expressing OSNs are limited to *D. suzukii* and *D. subpulchrella*, indicating that the change in their expression likely occurred in their common ancestor. However, because pseudogenised *Or67a.C1.1b* orthologs are still detectable in the genomes of the other Oriental Clade species and in *D. eugracillis* (Figure S2), we know that the origin of the gene predates the speciation events. This leaves open the possibility that *D. suzukii*’s and *D. subpulchrella*’s distinct *Or67a.C.1.1b* OSN expression may only appear to have originated recently due to multiple independent *Or67a.C1.1b* gene losses. Similarly, the new *Or67a.C1.1a*-expressing OSN can only be explained by multiple gains and/or losses. That *D. takahashii*’s *Or67a.C1.1a* is co-expressed in the ancestral ab10A-like OSNs, despite all other species with an intact ortholog having evolved the new OSN expression, highlights the dynamic expression changes that occurred for this receptor. Together, these findings provide the first evolutionary reconstruction of a transition of an *Or* “escaping” ancestral expression into a new OSN.

### Single-nuclei transcriptomics and *Or* pseudogenes provide resolution to cell type homology

The gain of expression in preexisting OSNs provides a parsimonious explanation for our *Or67a.C1.1a* and *Or67a.C1.1b* observations. This interpretation assumes, however, that the OSNs that we are calling ab2B and ab3A are homologous over the tens of millions of years that separate the species (Figure 1A). While considering this, we were reminded of the successful use of transcripts from *D. sechellia*’s *Or67a.C2* pseudogene for clarifying the evolutionary history of its *Or67a*-expressing OSNs (Figure 2C). We decided to test if a similar approach using the *Or22a* and *Or85a* pseudogenes would also provide clarification for ab2B and ab3A. If each of these OSNs are homologous, we would expect transcripts from *Or85a* and *Or67a.C1.1a* to colocalise in the same neurons, and we would expect the same for transcripts from *Or22a* and *Or67a.C1.1b*. Because it has the two new *Or67a*-expressing OSN populations, we again investigated this question in *D. suzukii* by generating and analysing a single nuclei RNA sequencing (snRNA-seq) dataset.

Using a set of 3 panneuronal and 4 OSN co-receptor marker genes, we subsetted the snRNA-seq dataset to neurons and further classified them into broad chemosensory clusters based on the expression patterns of *Orco, Ir8a*, *Ir25a*, and *Ir76b*, *Gr21a* and *Gr63a* (Figure 4A; Methods). We readily detected transcripts that mapped to *Or67a.C1.1a*, *Or67a.C1.1b*, and to a lesser extent *Or67a.C2*, but did not detect transcripts for *Or67a.C4*. Consistent with our HCR FISH results, *Or67a.C1.1a* and *Or67a.C1.1b* were found to be expressed in distinct olfactory neuron clusters (clusters 38 and 14+15, respectively), with faint *Or67a.C2* expression also detected in cluster 14+15 (Figure 4B,C). Somewhat surprisingly, we did not find an obvious ab10A cluster in our dataset, possibly due to lower nuclear transcript abundances for these receptors. Importantly, reads that mapped to the pseudogenised *Or22a* and *Or85a* loci were detected. Consistent with our expectations, these nuclei mapped to the same clusters as *Or67a.C1.1a* and *Or67a.C1.1b,* respectively, and to the same individual nuclei in multiple instances (Figure 4B). These results provide strong evidence that *Or67a.C1.1a* and *Or67a.C1.1b* have gained expression in ab2B and ab3A neurons, respectively, and further highlight the utility of using pseudogenes for reconstructing the evolutionary history of OSNs.

**Figure 4:**
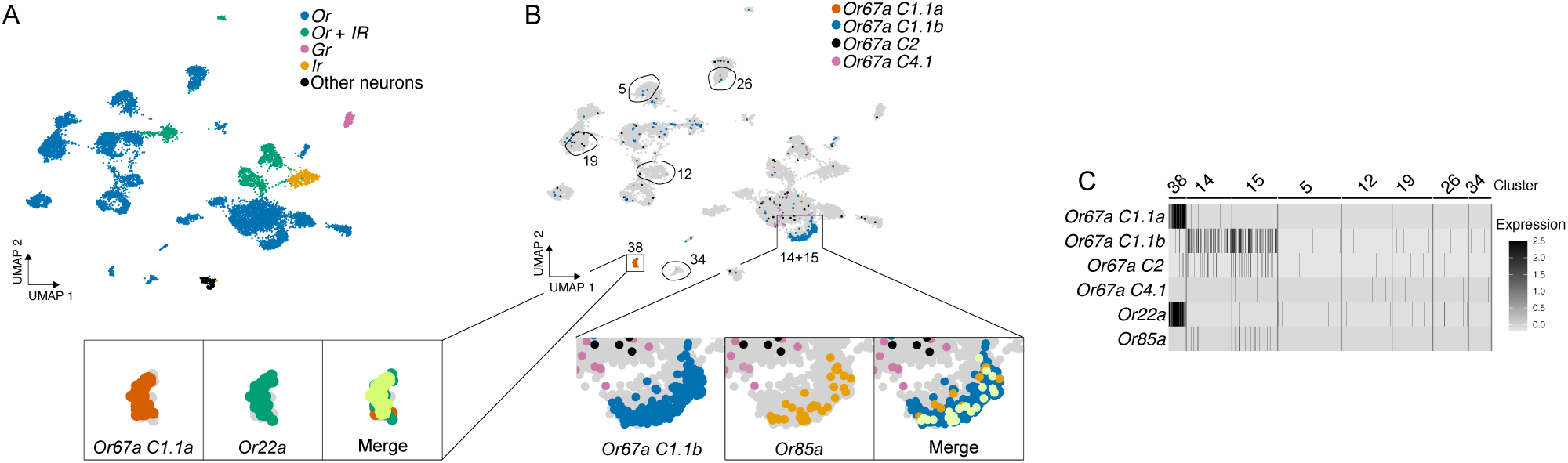
Co-expressed transcripts from pseudogenes resolved cell type homology. **A)** *D. suzukii*’s antennal snRNA-seq atlas showing only neuronal nuclei. Cell types are annotated according to the type of chemoreceptor that they express (*Gr* = Gustatory receptor, *Ir* = Ionotropic receptor, *Or* = Odorant receptor; Other = remaining neurons that were not found to express the former). **B)** Top: Atlas from panel A annotated with transcripts that map to *D. suzukii*’s four intact *Or67a* paralogs. Bottom: magnified view of clusters that express *Or67a.C1.1a* (38) and *Or67a.C1.1b* (14+15), along with nuclei that were found to contain transcripts from the receptors that they have replaced, *Or22a* and *Or85a*, respectively. **C)** Heatmap summarising the expression of the four *Or67a* paralogs and *Or22a* and *Or85a* from clusters 38 and 14+15 and five additional cell clusters chosen randomly (encircled in panel B).

### *Or67a*-expressing OSN populations have evolved in size and are sexually dimorphic

While carrying out the above HCR-FISH experiments, we noticed that the number of *Or67a*-expressing OSNs frequently varied within and between species. This led us to suspect that the size of the OSN populations could be sexually dimorphic and that their sizes may have evolved across species. To explore this further, we repeated most of the HCR-FISH experiments on an expanded and sex-separated set of antennae (Figure 5). This analysis indeed revealed significant sex differences across all *Or67a*-labelled OSN populations, with males having between 11-38% fewer OSNs compared to females (File S6). OSNs labelled by *Or67a.C1.1b* probes were the most dimorphic (rate ratio = 0.62, 95% CI 0.55–0.70; Wald *t*43 = −7.71; *p* = 1.2 × 10⁻^9^) while *Or67a.C3.1b* labelled OSNs were the least, though still significant, dimorphic population, (rate ratio = 0.89, 95% CI 0.80–0.98; Wald *t*43 = −2.29; *p* = 0.0269).

**Figure 5:**
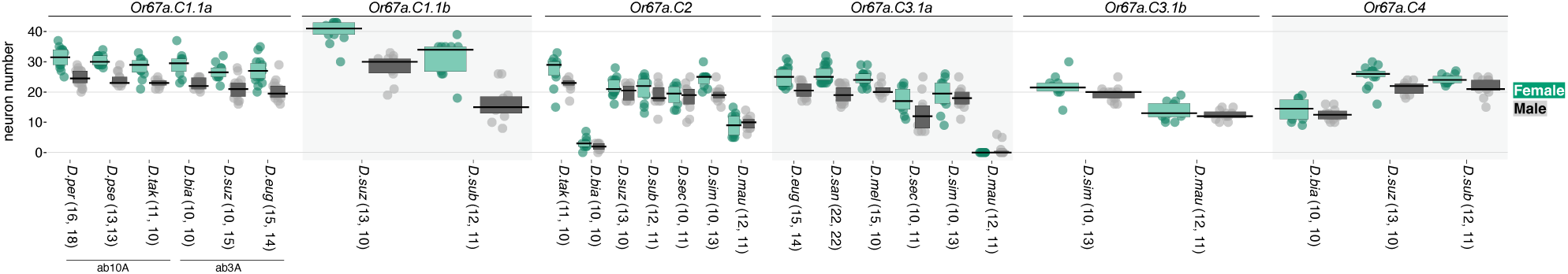
*Or67a*-expressing OSN are sexually dimorphic and evolve in cell number. Quantification of male and female OSN numbers across *Or67a* paralogs. Dots display replicate cell counts, rectangles span the interquartile range, and horizontal lines indicate median values. Species’ names are abbreviated to their first three letters. Numbers in parentheses are the sample sizes (number of antennae counted) for females and males, respectively.

We found that the average number of neurons labelled by the *Or67a* probes was ∼20 neurons, but varied significantly across species and all OSN populations, except for the ab3A neurons labelled by the *Or67a.C1.1a* probes (File S6). Among the *Or67a* copies that are co-expressed in the ab10A-like population (Figure 2C, D), we found that the neuron quantifications were largely consistent within species, suggesting uniform co-expression of the *Or67a* paralogs. The exceptions were in *D. mauritiana* and *D. biarmipies* (File S6). These inconsistencies in cell numbers may reflect subpopulations expressing different combinations of the *Or67a* copies and/or differences in labelling efficiency across probes. Excluding these latter three species from the analyses, significant species differences in the size of ab10A-like populations persist, however (File S6). These results show that, in addition to *Or67a.C1.1a* and *Or67a.C1.1b* having evolved new OSN expression, sexual dimorphism and population size differences across species are two additional modes of evolution of the *Or67a* subfamily. These results also add to the evidence that the population size of OSNs can evolve quickly^9,32–34^.

### Olfactory perception has diversified through *Or67a* neofunctionalization

The functional diversification of animal odorant receptors through gene duplication and neofunctionalization has been documented extensively^16,35–38^. Previously, we showed that adaptive changes that modified the odour responses of the co-expressed *Or67a.C2, Or67a.C3.1a,* and *Or67a.C3.1b* duplicates in *D. simulans* led to olfactory perception innovation^7^, and we wondered whether neofunctionalization has occurred more broadly throughout the *Or67a* subfamily. We were particularly interested in the possibility that *Or67a.C1.1a* and *Or67a.C1.1b* have evolved different receptor functions and thereby led to the evolutionary recoding of OSNs through the replacement of their ancestrally expressed receptors.

To address this question, we first used AlphaFold3^39^ to predict the paralogs’ structures in complex with three copies of their required co-receptor, Orco^40^ (using an intact *Or67a.C1.1a* allele). In each case, AlphaFold produced heteromeric assemblies with high confidence values, consistent with the formation of a functional Or67a–Orco complex (Figure S14). The predicted models adopt an open channel-like conformation, inferred from the displaced positioning of the gating helix, which in insect Ors is associated with ligand-bound, activated conformations. Although the overall architecture of the complexes is broadly conserved across the paralogues, the predictions reveal subtle but clear differences within the canonical ligand-binding region (Figure 6A-D). Consistent with this, P2Rank analysis^41^ identified binding pockets in all receptors at the expected canonical site, involving residues from helices 2, 3, 5, 6, and 8, together with contributions from the loop between helices 5 and 6 (Figure 6E). While the general pocket topology is maintained across the four receptors, the predicted pocket volumes differ. In particular, the Or67a.C1.1b model contains a substantially larger pocket than the other three Or67a paralogs, with a more open structure that incorporates additional residues from helix 7.

**Figure 6.**
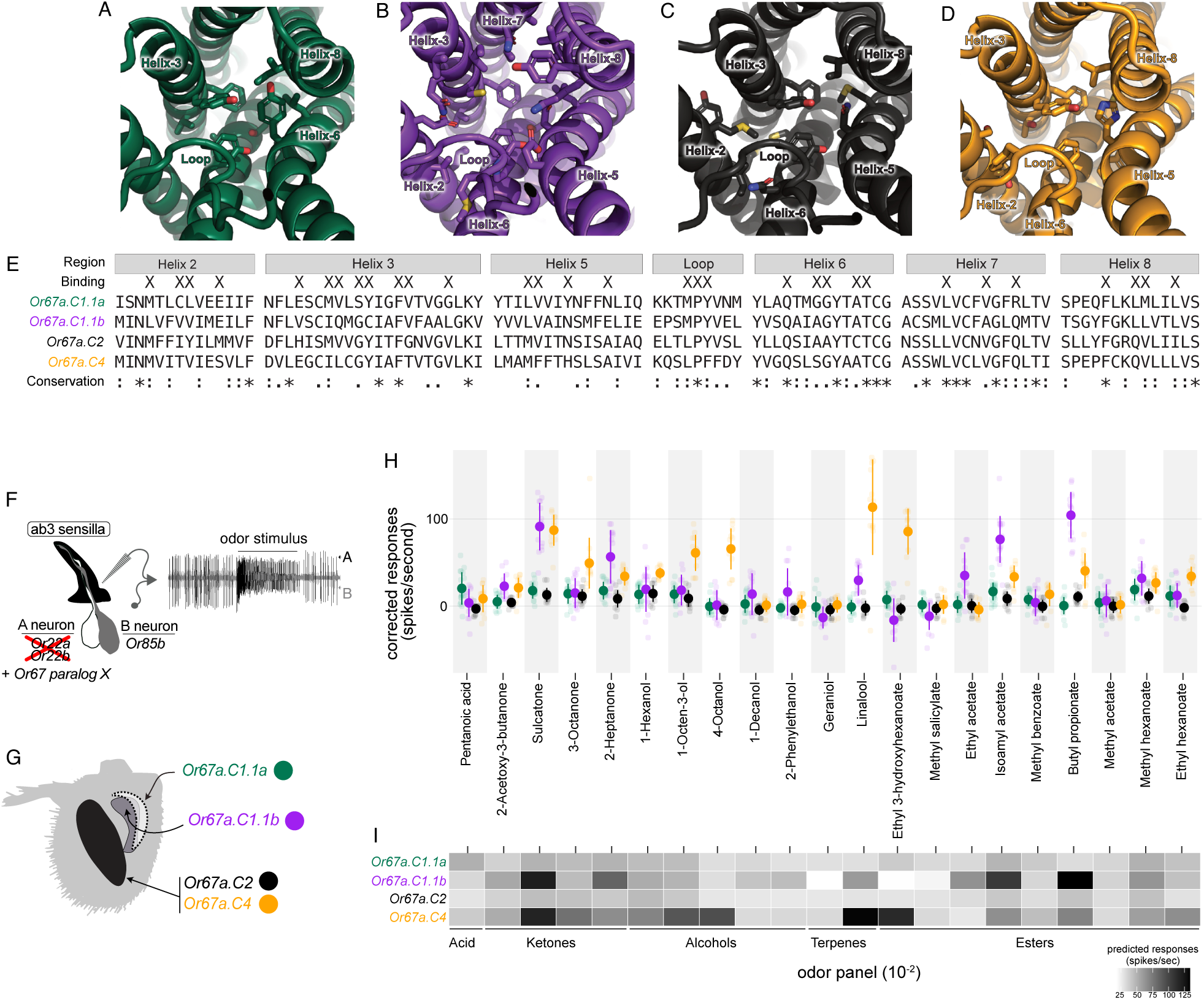
*Or67a* duplicates in *D. suzukii* have been neofunctionalised. **A-D)** Close-up view of the predicted binding sites across *D. suzukii*’s Or67a paralogs (see also Figure S14), coloured by receptor type Or67a.C1.1a (green), Or67a.C1.1b (purple), Or67a.C2 (black), and Or67a.C4 (orange). Side chains of key interacting residues from predicted binding pockets are depicted as sticks. **E)** Alignment of amino acid residues in the binding pockets, with an overview of positions in the structures based on helices shown, and with key interacting residues from P2RANK predicted binding pockets highlighted (“Binding” row). **F)** Schematic illustrating the decoder neuron system used to characterise *D. suzukii*’s Or67a paralogs. **G)** Schematic of an antenna and the OSNs that express the four Or67a paralogs characterised in the decoder system. Colors match panel H. **H)** Odour responses for Or67a paralogs across odour panel (each receptor-odor tested has between 11-18 replicates; all odour concentrations 10^−2^; transparent circles display the raw data; dark solid circles display the mean values and error bars display the standard deviation). **I)** Heatmap displaying the predicted responses based on modelling the odour response data using a GLM (Methods). Panel B’s x-axis is shared with the heatmap columns, with the category of compounds displayed below.

To experimentally test for functional differences, we performed *in vivo* electrophysiological recordings of odour-evoked activity. We individually expressed *D. suzukii*’s *Or67a.C1.1a*, *Or67a.C1.1b*, *Or67a.C2*, and *Or67a.C4* receptors in a *D. melanogaster* ‘decoder’ neuron – an ab3A neuron that lacks its endogenous tuning receptor (Or22a) but still expresses the required co-receptor, Orco^42^ (Figure 6F). Quantification of ab3A responses was made for a panel of 21 odours chosen as an expanded set of the classes of compounds that had previously been used to test *Or67a* paralogs^9^.

When preparing these experiments, we were surprised to find that *Or67a.C1.1a* contained a premature stop codon caused by a single-base deletion in a poly(A) tract of our lab strain (NGM-2; collected in Naganuma, Japan), suggesting that a nonfunctional allele is likely segregating at this locus. Alignment of the region across 7 available *D. suzukii* genome assemblies showed that the nonsense mutation is shared with one other strain (GB-ls-coga4, sampled from Bucharest, Romania; File S7). Because translational read-through has been reported for other *Drosophila Or* genes containing premature stop codons^43^, we included this allele in our experiment to test if a functional receptor is produced.

Analyses of the odour response data using a generalised linear model (Gaussian family, identity link) revealed receptors and odours had significant main effects, as did most of the receptor x odour interaction terms (File S8). Baseline responses were strongest for *Or67a.C1.1b* (β = 0.892, *p* < 2 × 10^−16^) and *Or67a.C4* (β = 0.435, *p* < 3.8 × 10^−15^), with each responding to multiple, and only partially overlapping, compounds (Figure 6G-I). *Or67a.C1.1b* responds most strongly to the esters butyl propionate and isoamyl acetate and to the ketone sulcatone. *Or67a.C4* also responded strongly to sulcatone, but its other strongest agonists were the ester ethyl 3-hydroxyhexanoate and the terpene linalool. *Or67a.C2* responded weakly to our odour panel, indicating that we did not include ecologically relevant compounds for it. *Or67a.C1.1a* also responded weakly to the panel, consistent with its early premature stop codon rendering it nonfunctional. In addition, its raw response traces contain very few of the signature large amplitude spikes that are expected when misexpressing functional *Or*s in the decoder neuron, furthering the case for it being a pseudogene in the NGM-2 strain (Figure S13).

During the completion of these analyses, an odour response dataset that also used the decoder neuron system for the four *Or67a* genes became available, providing an opportunity to qualitatively compare results across strains^44^. The comparison was especially meaningful for *Or67a.C1.1a*, because the published WT3 allele (sampled from California, USA) produced robust responses for 7 of 15 odours tested (>100 spikes/sec.), including 4 of 5 that overlap with our study (Isomamyl acetate, 2-Heptonone, 3-Octanone, and Methyl hexanoate)^44^. These observations mean that functional and nonfunctional *Or67a.C1.1a* copies are segregating in *D. suzukii*, and that several odour sensitives have been lost in ab3A neurons in the NGM-2 strain. Five odours also overlap between the *Or67a.C1.1b* experiments, for which good correspondence was found (Isoamyl acetate: ∼200^WT3^ vs. 104^NGM-2^, sulcatone: ∼200^WT3^ vs 121^NGM-2^, 2-heptanone: ∼70^WT3^ vs 85^NGM-2^, 3-octanone: ∼45^WT3^ vs 45^NGM-2^, and Methyl hexanoate: ∼40^WT3^ vs 62^NGM-2^ spikes/sec.). Only two odours were shared between the *Or67a.C2* and *Or67a.C4* studies, sulcatone and 1-Octen-3-ol, limiting their comparison: Both odour panels concordantly elicited weak *Or67a.C2* responses from the two strains (∼38^WT3^ vs. 45^NGM-2^ spikes/sec and ∼25^WT3^ vs. 40^NGM-2^ spikes/sec, respectively), but differences exist for *Or67a.C4,* with the *Or67a.C4*^NGM-2^ having stronger responses to both odours (∼20^WT3^ vs. 118^NGM-2^ and ∼30^WT3^ vs. 91^NGM-2^ spikes/sec, respectively).

These physiology results confirm the evolution of distinct odour tuning differences among *D. suzkukii*’s *Or67a* paralogs^44^. They also demonstrate a recoding of ab3A neurons through *Or67a.C1.1a*’s neofunctionalization and replacement of *Or22a* and a parallel recoding of ab2B neurons through *Or67a.C1.1b*’s neofunctionalization and replacement of *Or85a*. That *Or67a.C1.1a* segregates a polymorphic null allele further highlights the continuous changes that the receptor and its respective OSN population experiences. Finally, the response differences between the *Or67a.C2* and *Or67a.C4* suggest that *D. suzukii*’s ab10A OSN are tuned through the co-expression of distinct Ors. OSN tuning by co-expressed Ors would be analogous to our previous observations in *D. simulans*^7^, though knockouts of the two receptors and quantification of ab10A odour responses are needed for verification.

## Discussion

We uncovered a spectrum of *Or* duplicate fates that link *Or* copy number changes with the diversification of olfactory perception. We found that co-expression is the most likely initial phase of an *Or* duplicate. As expected under a birth-and-death model of gene duplication, many of these co-expressed *Or*s become pseudogenised through the accumulation of disruptive mutations^45,46^ (Figures S2-10). But for those *Or* duplicates that remain intact and fix within a species, we showed that co-expression of functionally divergent duplicates can provide a stable and, in some cases, adaptive path for neurons to gain novel odour response properties^7^. With the discovery of *Or67a*.*C1.1a’s* and *Or67a*.*C1.1b’s* new expression patterns, we identified and reconstructed an additional fate for *Or* duplicates in which regulatory changes place the receptors in different but pre-existing OSN populations. These results demonstrate that expansions in the *Or* gene family that result in new perceptual changes do not require the evolution of new OSN lineages.

As additional cross-species studies are carried out, we anticipate that analogous changes will be identified for other *Or* subfamilies (and duplicates within other chemoreceptor families^e.g.,47^). The combination of *Or* protein and neuron-specifying regulatory changes may provide flexible routes for altering peripheral nervous systems, and potentially for behavioural valence changes depending on the context of the olfactory circuit in which a receptor gains new expression. Specific to the *Or67a* subfamily, further case-studies within the Oriental Clade, along with population-level analyses, will help us understand if the ancestral *Or* losses that we observed for ab3A (*Or22a*) and ab2B (*Or85a*) were coincidental. The inclusion of additional species will also help resolve the number of times *Or67a*.*C1.1a* and *Or67a*.*C1.1b* have gained new OSN expression, and will be key for identifying the genetic bases of their new expression.

*D. suzukii* females are unusual in their attraction to and ovipositing in ripe fruit. Their larvae destroy the fruit as they develop, resulting in costly damage to the fruit industry^48,49^. Because *D. suzukii* has recently expanded globally from East Asia, there is increasing interest in understanding the unique features of its chemosensory systems, particularly from a management perspective^50^. Remarkably, the expression and functional innovations involving *Or67a.C1.1a* and *Or67a.C1.1b* provide the molecular and neuronal bases of two of the most pronounced olfactory differences that were previously identified in a systematic physiological screen that compared *D. suzukii*’s antennal responses to *D. melanogaster*’s^31^. Though most odour responses were conserved between the two species, the screen - which was blind to the receptors being activated - identified ab2B and ab3A neurons among the few showing differences. Chemical ecology studies have suggested that ab2B’s sensitivity shift towards ripe fruit compounds such as isoamyl acetate (compared to overripe compounds), combined with ab3A’s shift to include compounds such as β-cyclocitral, which are associated with leaf tissues, may have been key sensory modifications^31,44,51^. Speculations about the causes of the two OSN differences in *D. suzukii* had previously been made^31^, but only recently was *Or67a.C1.1b* implicated in the ab2B changes through additional *in vivo* electrophysiological experiments and an investigation of odour-evoked responses that were lost when *Or67a.C1.1b* was knocked out^44^. Here, we provided direct evidence that *Or67a.C1.1b* is expressed in a new neuron population and that the population is homologous to *D. melanogaster*’s ab2B OSNs. We’ve also shown that OSN recoding events like this one is repeatable by discovering a parallel event involving *Or67a.C1.1a* and the ab3A neuron. Our evolutionary and functional analyses have reconstructed the transitional steps underlying the recoding of preexisting OSNs by duplicated *Or*s and have experimentally linked these changes to species-specific sensory differences.

## Methods

### Genes identification and annotations

The *Or67a* copies were retrieved in the following 18 drosophila species annotated genomes: *D. biarmipes* (GCF_018148935.1), *D. erecta* (GCF_003286155.1), *D. eugracilis* (GCF_018153835.1), *D. lutescens* (home-made), *D. mauritiana* (GCF_004382145.1), *D. melanogaster* (GCA_000001215.4), *D. orena* (GCA_005876975), *D. persimilis* (GCA_020698555.1), *D. pseudo-obscura* (GCA_009870125.2), *D. santomea* (GCA_016746245.2), *D. sechellia* (GCA_004382195.1), *D. simulans* (GCF_016746395.1), *D. subpulchrella* (GCA_014743375.1), *D. suzukii* (GCF_043229965.1), *D. takahashii* (GCA_030179915.1), *D. teissieri* (GCA_016746235), *D. yakuba* (GCA_016746365), *D. virilis* (GCA_007989325.2). *D. melanogaster* and *D. suzukii* Or67a protein sequences were first used as queries in TBLASTN against available genomes on the NCBI database. Nucleotide sequences of the *Or67a* copies obtained through that process were then aligned on the genomes detailed above using Minimap2^52^ (v2.17) within Geneious (v2022.22.2). All sequences found subsequently were used for a second round of BLAST and alignment using BLASTN, and Minimap across the above genomes to identify copies that may have been missed in the first round. The sequences obtained from these iterations were aligned using Clustal Omega^53^. Coding sequences were generated using pre-existing annotations of intronic sequences and by manual inspection. The coding sequences were translated into protein using Geneious. Genomic, coding and protein sequences used for subsequent analysis are available in File S1.

### Phylogenetic analysis

Protein sequences were aligned using Clustal Omega within Geneious (v2022.22.2) using default settings. To infer the Or67a family protein tree, we included 53/56 protein sequences (the *Dbia.C1.1b*, *Deug.C1.1b*, *Dtak.C1.1d* were excluded) and used Mr.Bayes (v3.2.6)^54^ with the following settings: prset aamodelpr=mixed; mcmc nchains=4; nruns=2; ngen=20000; samplefreq=100 printfreq=100 diagnfreq=1000 Temp=.03; sump burnin=5000; sumt burnin=5000. Dates provided for Figure 1A’s tree were sourced from ^29,55–57^.

### Microsynteny analyses

The microsynteny of *Or67a*-containing regions was investigated across species by extracting 20 megabases (Mb) upstream and downstream of each *Or67a* copy identified through the homology search described above. These 40 Mb intervals were then iteratively aligned to the other species’ genomes using Minimap2 (v2.17) in Geneious to refine the start and end positions of the homologous regions. The final set of orthologous *Or67a*-containing intervals was aligned using MAFFT^58^ (v1.5.0). For non-*D. melanogaster* species, we annotated the genes flanking *Or67a* copies within these intervals using NCBI’s RefSeq genome annotations when available. For *D. melanogaster* annotations, FlyBase (FB2023_05 September) was used. We additionally verified that the flanking genes are orthologous using BLAST+^59^. The chromosomal locations of the Or67a-containing intervals were determined from genomes with chromosome-level assemblies (D. melanogaster, D. yakuba, D. simulans, D. santomea, D. teissieri, D. takahashii, and D. pseudo-obscura) and those with near-chromosome-level assemblies (*D. suzukii* and *D. subpuchrella*). Alignments using the aforementioned genomes allowed us to identify the chromosome location for the intervals within *D. biarmipes*, *D. erecta*, *D. eugracilis*, *D. lutescens*, *D. mauritiana*, *D. orena*, *D. persimilis*, *D. sechellia*, which have less complete assemblies. Transposable elements within the intervals were annotated using the *Drosophila* repbase dataset^60^. TEs and genes were manually annotated within homologous regions in Geneious. R’s gggenomes visualisation package^61^ (v1.1.3) was used to generate the microsynthenic alignment plots (Figures S2-10). Microsynthenic alignments for each clade are available in File S2.

### Test for TE enrichment

*D. suzukii*’s and *D. takahashii*’s *Or67a*-containing intervals were found to contain the most TE fragments (Figures S2-10). To test for an association between their *Or67a* genes and TE sequences, we generated a null distribution of nucleotide overlap by repeatedly sampling (n=5000) the same number of genes of approximately the same length (800-2500bp) from their annotated genomes (*D. suzukii*: Dsuz_2.1.pri and *D. takahashii*: DtakHiC1). We treated the tandem duplicates, *Or67a.C1.1a* and *Or67a.C1.1b*, of both species as a single locus. We then computed the fraction of nucleotides that overlapped annotated TEs using BEDtools (v2.31.1)^62^ and compared this distribution to our empirical value (see TE_enrichment in our GitLab Repository: https://gitlab.com/EvoNeuro/Or67a_NewPops).

### Paml analysis

To quantify the selective constraints acting on *Or67a* copies, a codon-based likelihood method was applied within the CODEML program (v.4.8)^63^ implemented the pamlX GUI (v.1.3.1)^64^. The 46 non-pseudogenized *Or67a* copies were aligned using the translation alignment function in Geneious (v2022.2.2), with gap-containing sites manually removed where necessary. The resulting alignment, along with the species topology, was used to infer a gene tree using Mr.Bayes (v3.2.6)^54^, with the same settings described above. We used CODEML’s branch models to test whether specific lineages or groups of lineages evolved under different selective regimes than the rest of the protein tree^65^. For each test, the branches of interest were designated as foreground, while all remaining branches constituted the background. Branch groups were defined based on phylogenetic relatedness and taxonomic groupings. We first estimated a single ω across all branches (model = 0, NSsites = 0) as a null model. We then allowed the foreground and background branches to have distinct ω values (model = 2, NSsites = 0), and in some cases tested models with three or more ω classes to further partition selective regimes among subgroups. Log-likelihood values of nested models were compared using a likelihood ratio test (LRT), with test statistics approximated by a χ² distribution with degrees of freedom equal to the difference in the number of parameters between models. Each test addressed an independent biological hypothesis regarding a specific lineage or clade; therefore, no correction for multiple testing was applied. Results for all models tested are provided in File S3.

### Fly rearing

Fly strains were reared on a standard yeast/cornmeal/agar medium supplemented with Carolina 4-24 Formula (Carolina Biological Supply) and maintained in a 12:12 hr light: dark cycle at 25 degrees. Adults between 2 to 9 days old were sorted and sexed on CO2 at least 24h before the dissections or physiology experiments.

### HCR RNA fluorescence in situ hybridization and immunofluorescence

RNA in situ hybridization reactions were performed as described previously^66^ on *D. eugracilis* E-18101, *D. pseudo-obscura* k-s12, *D. persimilis* k-s11, *D. simulans* 14021-0251.008, *D. sechellia* 14021-0271.07, D. mauritiana MS9, *D. santomea* 14021-0271.00, *D. subpulchrella NGN5*, *D. suzukii* NGM-2, *D. biarmipes* 14023-0361.03, D. takahashii IHYT1. Flies aged between 2-10 days were flash-frozen in liquid nitrogen and agitated over a mini-sieve connected to a collection dish. Antennas were recovered from the collection dish using a pipette under a dissecting scope. They were collected on PBT (1X PBS, 0.1% Triton X-100) and fixed in 2ml of a 4% paraformaldehyde, 1X PBS, 0.1% Triton X-100 solution at 4 °C on a rotator set to low speed (<20 rpm) for 2 h. Following fixation, samples were washed twice in PBS + 3 % Triton X-100 and three times in PBT. The protocol suggested by Molecular Instruments for generic samples in solution was then followed (https://files.molecularinstruments.com/MI-Protocol-RNAFISH-GenericSolution-Rev9.pdf). Following the washes, samples were mounted in Vectashield and stored at 4°C. HCR probes set, amplifiers, and buffers were purchased from Molecular Instruments. Probe sets were designed by Molecular Instruments to maximize hybridization efficiency while minimizing off-target complementarity. Given the high sequence homology among the target genes, particular attention was paid to specificity during the design process. The specificity of each probe set was verified by Molecular Instruments during design and subsequently confirmed by the authors. While probe sequences are proprietary to Molecular Instruments and cannot be disclosed, the target sequences used for probe design, the species in which each probe set was applied, and the amplifiers used are listed in File S9.

Co-expression patterns of *D. yakuba*’s *Or67a*.*C3.1a* and *Or67a*.*C3.1b* copes were imaged using a transgenic driver line *Or67a*.*C3.1a* (w-; *Dyak*.*Or67a*.*C3.1a-RFP, w+*, see pDEST constructions, below) together with a standard FISH probe for *Or67a*.*C3.1b*. Flies were aged between 2-10 days and flash-frozen in liquid nitrogen then agitated over a mini-sieve connected to a collection dish. Antennas were recovered from the collection dish using a pipette under a dissecting scope. They were collected on PBT (1X PBS, 0.1% Triton X-100) and fixed in 2ml of a 4% paraformaldehyde, 1X PBS, 0.1% Triton X-100 solution at 4 °C on a rotator set to low speed (<20 rpm) for 2 h. Following fixation, samples were immersed for 20 min in methanol, then stepwise rehydrated (75%, 50%, 25% methanol in PBT, followed by two washes in PBT). Samples were then permeabilised with proteinase K (1 μL/mL in PBT) for 15 min at room temperature. Hybridisation was performed using a DIG-labeled probe specific to Dyak.*Or67a*.*C3.1b* diluted in hybridisation buffer and incubated at 60°C for 72 h. Subsequent steps were performed as previously described (Saina and Benton, 2013). Following the washes, samples were mounted in Vectashield and stored at 4°C.

Expression of *Dmel.Or67a.C3.1a* in antenna was examined using the *Dmel.Or67a.C3.1a–mCD8::GFP* reporter line^30^. Antennae were dissected and fixed as described above. Following three 10 min washes in PBT and two 10 min washes in SSCT, the residual liquid was removed, and the antennae were mounted in Vectashield and stored at 4°C.

Generalized linear models were used to analyze the effects of species and sex on the number of cells labeled by the *Or67a* HCR FISH probes (or, for *D. melanogaster*, based on the Dmel.Or67a.C3.1a–mCD8::GFP reporter line). The number of cells was fitted with a quasipoisson error distribution and a log link in R (v4.4.1) with the glm() function, with fixed effects set as species and sex (count ∼ species + sex). Model fit and dispersion were assessed using residual deviance and Pearson residuals. We also evaluated the impact that unequal variance might have on our model’s standard errors by applying a robust covariance estimator using the vcovHC() function in the sandwich library (v3.1-1)^67^. Significance of coefficients was evaluated using t-statistics derived from the quasi-likelihood framework. Replicates are independently stained antennae (n = 10-22 antennae per sex/species or 22-44 per species). Full model output is provided in File S1 and analysis code and results are available in the EvoSizeXSexSpecies.html.Rmd file within our GitLab repository: https://gitlab.com/EvoNeuro/Or67a_NewPops.

### Image acquisition

Antenna images were acquired on inverted confocal microscopes (Zeiss LSM 710 or LSM 880 or Leica Stellaris 5 or Leica Stellaris 8) equipped with an oil immersion 40X objective (Plan Neofluar 40X oil immersion DIC objective; 1.3 NA or HC PL APO CS2 40X oil immersion; 1.3 NA). The images were processed in Fiji (v1.53^68^). OSN numbers were counted using the Cell Counter Plugin in Fiji.

### Generation of transgenic lines

#### pUAST-attB

*DsuzOr67a.C1.1a, DsuzOr67a.C1.1b*, *DsuzOr67a.C2* and *DsuzOr67a.C4* coding sequences were amplified by PCR from *D. suzukii* NGM-2 cDNA. *Dsuz.Or67a.C1.1a* and *Dsuz.Or67a.C1.1b* were digested by XbaI/EcoRI and *Dsuz.Or67a.C4* by XbaI/BglII, (New England Biolabs) and inserted into a linearized pUAST–attB^69^. *DsuzOr67a.C2* coding sequence was first cloned into a pGEMTe vector (Promega), then amplified by PCR from the plasmid and integrated into an EcoRI/XbaI linearized pUAST–attB through Gibson assembly using the ClonExpress Ultra One Step Cloning Kit (Vazyme). Injections were performed in *D. melanogaster vas-int; attP-86Fb* flies (BDSC no. <u>24749</u>) by Rainbow Transgenic Flies for pUASt-attB-Dsuz.Or67a.C1.1b and by the Cambridge’s *Drosophila* Microinjection Services for pUASt-attB-DsuzOr67a.C1.1a, pUASt-attB-Dsuz.Or67aC2 and pUASt-attB-DsuzOr67a.C4.

#### Or67a reporter vectors: pDEST constructions

Predicted promoter region of *D. yakOr67a.C3.1a* was amplified by PCR from genomic DNA and inserted into pENTR/D-TOPO vectors (Thermo Fisher Scientific). The resulting vectors were then recombined with pDEST–HemmarR^70^ destination vectors via an LR recombination reaction (Gateway, Thermo Fisher Scientific). Transgenesis was performed in *D. yakuba#1734*^71^ by Rainbow Transgenic Flies. Positive transformants were crossed with *D. yakuba* w- flies (gifts from D. Stern).

#### Single-nuclei RNA-seq data generation

10x Antennal scRNA-sequencing was performed on *D. suzukii* NGM-2 partially following the protocol established by the Liqun Luo lab. Flies aged between 2-10 days were flash-frozen in liquid nitrogen and agitated over a mini-sieve connected to a collection dish. Two hundred antennas were collected from the collection dish using a pipette under a dissecting scope. They were transferred to 1.5 ml Eppendorf tubes containing 100 μl of PBS and stored at −80°C. 100 ul of the homogenization buffer was added to the samples, which were ground for 30 seconds with a pellet pestle motor. After the addition of 900ul of homogenization buffer, the samples were transferred into a dounce, and the nuclei were released by 30 strokes of the loose pestle, and 30 of the tight, on ice. Samples were filtered twice, once with a 5ml cell stainer and then on a 40um Flowmi. They were centrifuged 10min at 1000g at 4°C. The supernatant was discarded and the pellet resuspended in 1ml of PBS+0,5%BSA, filtered on a flowmi, stained with Hoechst 33342 and loaded on a MoFlo Astrios EQ cell sorter. The quality of the samples and the number of cells per sample were assessed using a Luna FX cell counter. Around 10000 cells were sorted per sample. Between 2 or 3 Samples were loaded onto a Chromium Next GEM Chip (10x Genomics). The GTF facility operated the 10X Genomics Chromium, generated the sequencing library and performed the 10X library sequencing on a NovaSeq.

#### Single-nuclei RNA-seq analyses

##### Updating reference annotations

Manual updates were made to the mitochondrial genome and *Or67a* copies in *D. suzukii*: mitochondrial annotations from the *D. melanogaster* reference genome (GCF_000001215.4) were blasted (BLASTN vX (REF)) against the *D. suzukii* genome (GCF_013340165.1). Any previously unannotated genes were added to the GTF file later used in mapping. The updated GTF file was then used with its corresponding genome to map reads using the STAR aligner (v2.7.11b^72^). A genome index was created with the ‘genomeGenerate’ mode and then reads were mapped using the ‘alignReads’ mode. To generate the final GTF file, the BAM file from the initial alignment was used with the tool peaks2utr (v1.4.1^73^) to annotate/extend 3’ UTRs with [--min-pileups 5 –extend-utr –do-pseudo --gtf] flags specified (the remaining settings were left as default).

##### Mapping sequencing reads, filtering, and integration

The updated annotations were once again used with the above *D. suzukii* genome and STAR aligner (v2.7.11b^72^) to generate a genome index with the command ‘genomeGenerate’ and then map sequencing reads using the command ‘alignReads’ The outputted count matrices were used for all subsequent analysis in RStudio^74^ (v2025.9.2.418). Steps prior to merging were completed in parallel for both biological replicates. Using a custom function (‘filter_droplet’; see GitLab repo, below) we imported the ‘full’ and ‘exon’ raw count matrices, and filtered out droplets with fewer than 70 UMIs or an intronic proportion ≤0.09. The SoupX^75^ package (v1.6.2) was run with both the ‘full’ raw count matrix before and after intronic proportion filtering as inputs to remove ambient RNA contamination. This count matrix was then used to create a Seurat object^76^ (v5.2.1). Mitochondrial and ribosomal gene percentages were used as a measure of effective nuclei separation, and droplets with a mitochondrial percentage >3.0 or a ribosomal percentage >10.0 were filtered out. Mitochondrial and ribosomal genes were removed to prevent their usage in dimensionality reduction steps. Droplets with <250 UMIs or <150 genes were also filtered out. The filtered Seurat object then underwent normalisation and dimensionality reduction in line with DoubletFinder’s^77^ (v2.0.4) recommended practice. DoubletFinder was run with an adjusted 5% doublet expectation (based on number of sequenced nuclei). Any remaining nuclei with >5500 UMIs or >2200 genes were also removed. Biological replicates were assessed for consistency by comparing number of cells, the total number of genes, the number of clusters, and the local topology of transcriptomic atlases. Replicate objects were merged and integrated using Seurat’s ‘IntegrateLayers’ function with the CCA Integration option.

##### Neuronal subsetting and cluster annotation

Marker expression was quantified using UCell’s ‘AddModuleScore_UCell’ function^78^ (v2.8.0). Nuclei with a UCell score ≥ 0.25 for panneuronal markers (*bruchpilot, Synaptotagmin 1, embryonic lethal abnormal vision*) or for OSN co-receptors *Orco, Ir8a, Ir25a* or *Ir76b* were selected for the neuronal subset. This subset underwent normalisation and dimensionality reduction again. Clusters were then annotated with an OSN type based on the following markers: *Or*: *Orco*, *Ir*s: *Ir8a, Ir25a, Ir76b*, *Gr*s:*Gr21a, Gr63a*, or ‘other neuronal’ for clusters where these markers were not found. Expression of the four intact *D. suzukii Or67a* paralogues (*Or67a.C1.1a*, *Or67a.*C1.1b, *Or67a.*C2 and *Or67a.*C4) and those co-express with (*Or22a* and *Or85a*) were plotted from this dataset.

Code for our snRNA-seq analyses can be found in our lab’s repository: https://gitlab.com/EvoNeuro/Or67a_NewPops/. Sequence Read Archive accession IDs for the raw snRNA-seq data will be made available upon publication.

#### Protein modelling

Protein Data Bank accession codes for all PDB files will be provided upon acceptance.

#### Electrophysiology

The *D. melanogaster* “decoder neuron system”^79–81^ was used to characterize Or67a paralog odour responses *in vivo*. Transgenic UAS lines (see pUAST-attB section, above) were crossed to *D. melanogaster* Δhalo/Cyo; Or22A-Gal4 flies. The resulting progeny expressed the *D. suzukii* Or67a paralogs in ab3A olfactory sensory neurons (OSNs) that lack the endogenous Or22a receptor. Single-sensillum recordings were performed as described previously^82^ using an odour concentration of 10^−2^. All odours were diluted in paraffin oil, except for pentanoic acid, 2-Acetoxy-3-butanone and 1-hexanol, which were diluted in double-distilled water. All flies used in these experiments were 3–10 days post-eclosion. Odorants and their CAS numbers are listed in File S9.

Odours were presented in 500 ms pulses using a Syntech CS-55 stimulus controller, with a minimum inter-stimulus interval of 60 s. Corrected responses were calculated as the number of spikes in a 500 ms window during odour stimulation (3.2–3.7s after recording onset), minus the number of spontaneous spikes in a 500 ms pre-stimulus window (1.2–1.7 s before stimulus onset). The spike count was then multiplied by two to obtain spikes per second. Solvent-corrected responses were calculated by subtracting from the response to each diluted odour the response obtained when stimulating with the corresponding solvent. Recordings were performed on a maximum of three sensilla per fly. Spike visualisation and quantification were performed using Spike2 (CED).

A generalized linear model (GLM, Gaussian family and identity link) was used to quantify how Or67a paralog, odor identity, and their interaction shape odour evoked responses. The analyses were performed in R (version 4.5.1; glm function in the stats package). Corrected spike counts were transformed by subtracting the minimum response and taking the natural logarithm. Model parameters were estimated using maximum likelihood under Fisher scoring iterations (2 iterations to convergence). Model selection without (resp_log ∼ receptor + odor) and with (resp_log ∼ receptor + odor + receptor:odor) an interaction term was assessed using the Akaike’s Information Criterion (AIC). AIC showed an substantial improvement for the model with the interaction term over the main effects model (ΔAIC = 734.72) and was kept for subsequent analyses. No regularization, random effects, or post hoc corrections were applied. All analyses were performed on 1259 observations after removing three outliers (three extreme negative *Or67a.C1.1b* responses to 1-Hexanol, Ethyl 3-hydroxyhexanoatem, Methyl salicylate). Code for these analyses can be found in our GitLab repository: https://gitlab.com/EvoNeuro/Or67a_NewPops/.

## Supporting information

SupFiles 1-9

## Data Availability

All data generated or analysed during this study are included in this published article, its supplementary information files, and GitLab repro (https://gitlab.com/EvoNeuro/Or67a_NewPops/)

## Acknowledgements

We thank members of the Arguello lab and Margarida Cardoso-Moreira for insightful feedback throughout the course of this work; Margarida Cardoso-Moreira, Thomas O. Auer, Juan Antonio Sánchez-Alcañiz, and Juan Lugo Ramos for comments on earlier drafts of this manuscript; Richard Benton for temporary use of his SSR rig; QMUL’s Apocrita HPC and Swiss Institute of Bioinformatics for providing computational resources; T.M. Balls for provided support and inspiration; Research in JRA’s laboratory was supported by Queen Mary University of London, the University of Lausanne, and the Swiss National Science Foundation (grants PP00P3_176956 and 310030_201188).

## Author contributions

Conceptualisation: GB, JRA

Writing - original draft: GB, JRA

Writing - review & editing: GB, SH, JRA

Methodology: GB, TB, SH, TK, BM, VV, MD, JRA

Investigation: GB, TB, SH, MD, JRA

Visualisation: GB, SH, MD, JRA

Funding acquisition: JRA

Project administration: GB, JRA

Supervision: MD, JRA

## Supplementary Files

File S1: All Or67a sequences (DNA and protein) used in the study in fasta format

File S2: Syntenic alignments for the Or67a-containing regions in fasta format.

File S3: Table containing PAML results.

File S4: Or22a protein sequence alignments in fasta format.

File S5: Or85a protein sequence alignments in fasta format.

File S6: Data and results for Or67a counts across species and sexes.

- Sheet 1: Raw count data
- Sheet 2: Overall sample size summary for the dataset
- Sheet 3: Sample size per species and per sex
- Sheet 4: Sex-averaged number of Or67a-expressing OSN across species
- Sheet 5: GLM result on full dataset
- Sheets 6-17: GLM applied to each Or67a paralog across species. Each analysis has two tables, one with the coefficients and one with the coefficients translated into cell numbers.
- Sheet 18: Comparison of average counts for co-expressed Or67a copies

File S7: FASTA alignment of poly(A) tract showing the single base deletion causing a premature stop codon in the NGM-2 strain.

File S8: GLM results for the SSR data.

File S9: Table containing reagents used in the study.

---

**Figure S1:**
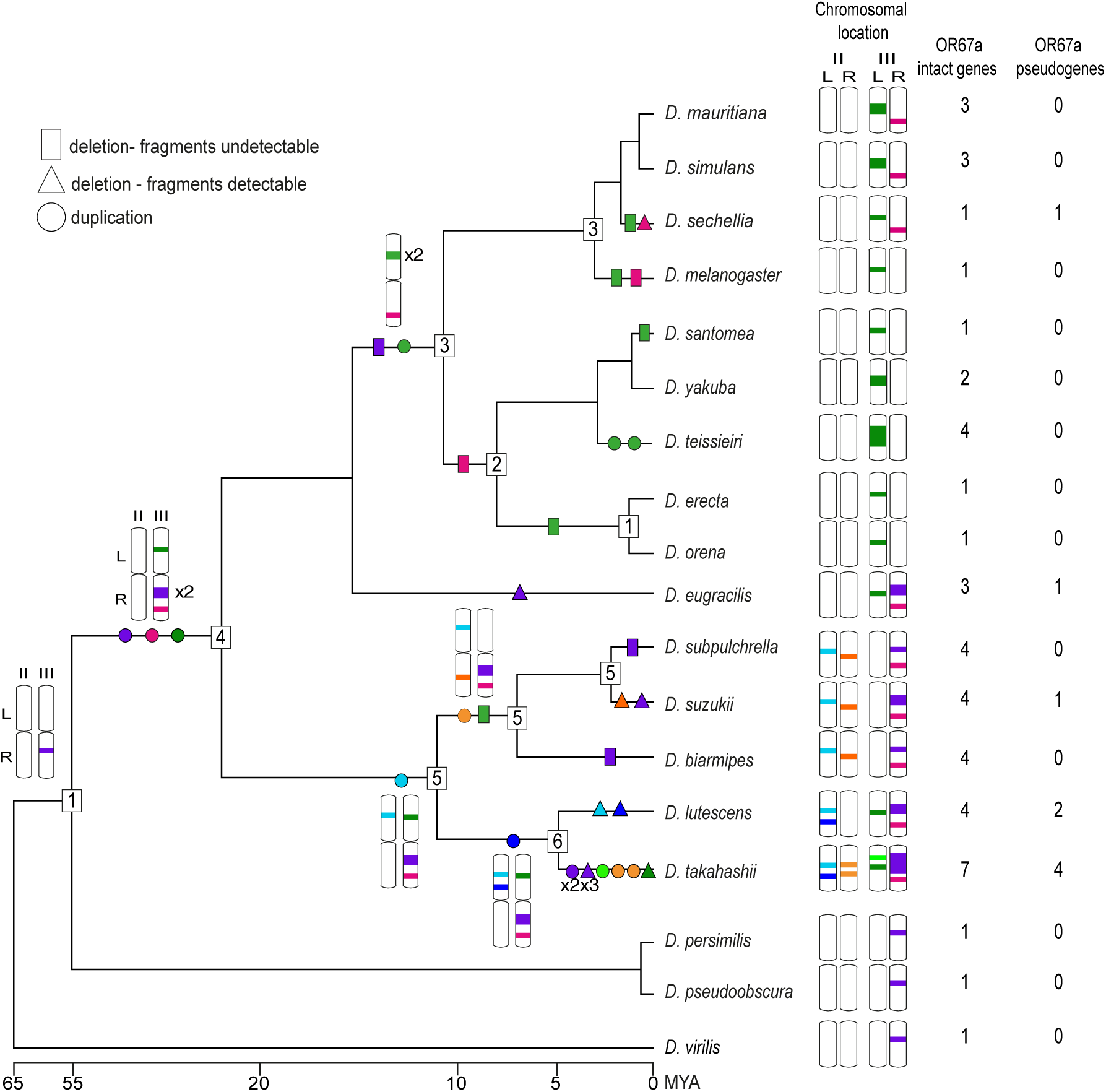
Evolution of the *Or67a* subfamily within the *Drosophila melanogaster* species group. Number and chromosomal locations of intact and pseudogenised *Or67a* paralogs. The number of genes inferred at ancestral nodes is indicated in boxes. Chromosomal locations of *Or67a* paralogs are shown at the tree tips, along with the number of intact and pseudogenised copies.

**Figure S2:**
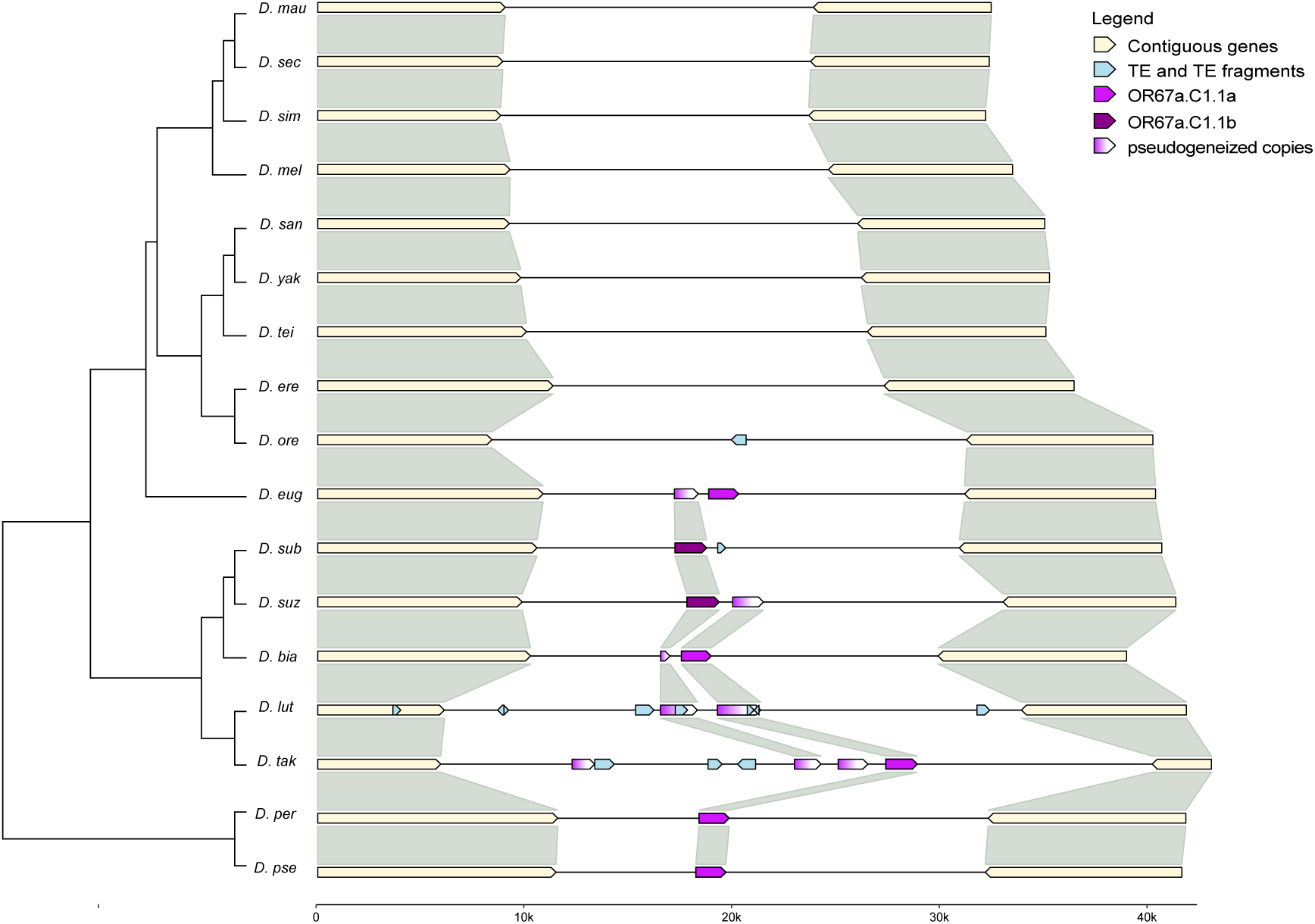
Microsynteny analysis of the *Or67.C1.1* cluster. Alignments are arranged by phylogenetic relationships (tree in left margin, not to scale). Species names are abbreviated using the first four letters of the genus and species.

**Figure S3:**
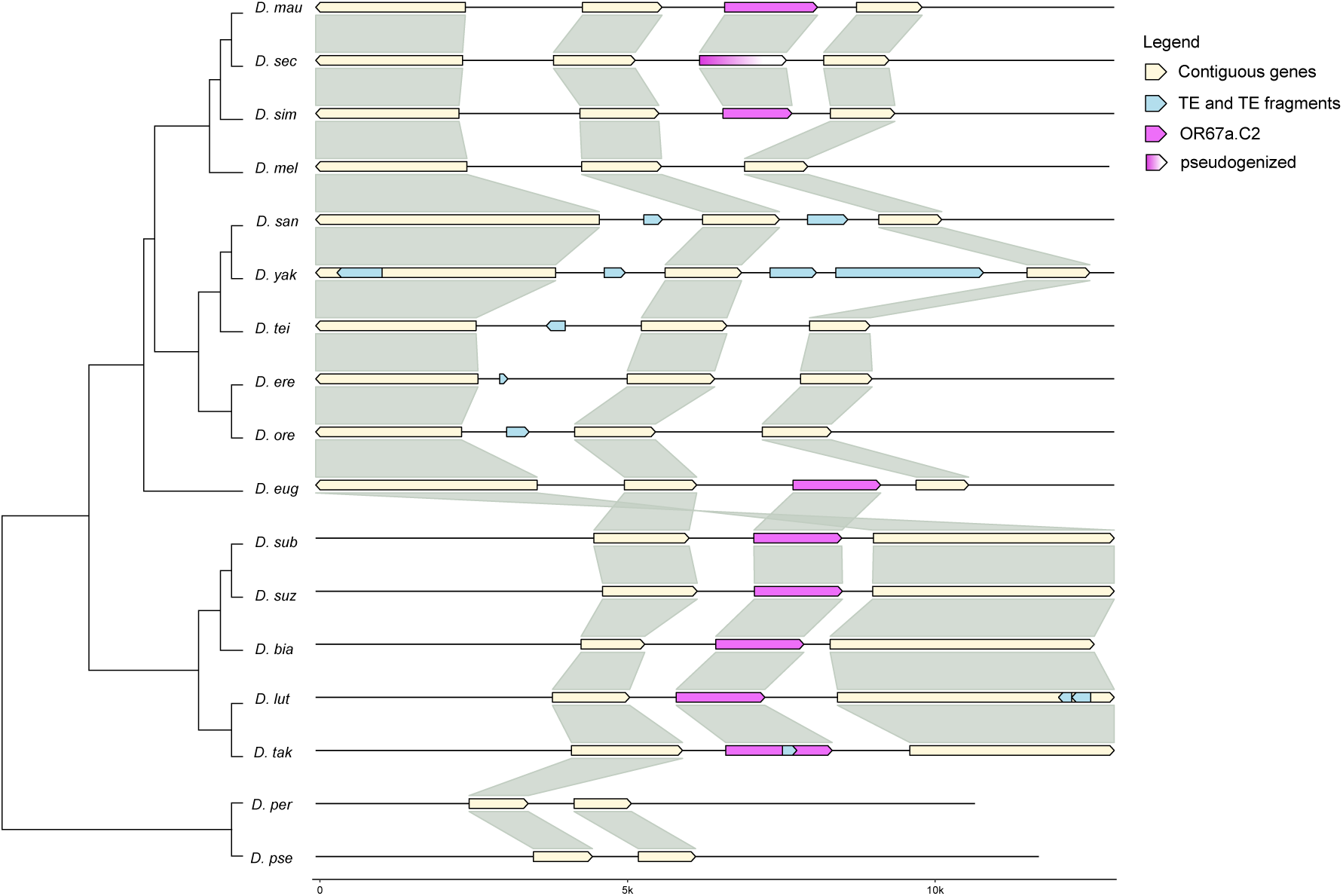
Microsynteny analysis of the *Or67.C2* cluster. Alignments are arranged by phylogenetic relationships (tree in left margin, not to scale). Species names are abbreviated using the first four letters of the genus and species.

**Figure S4:**
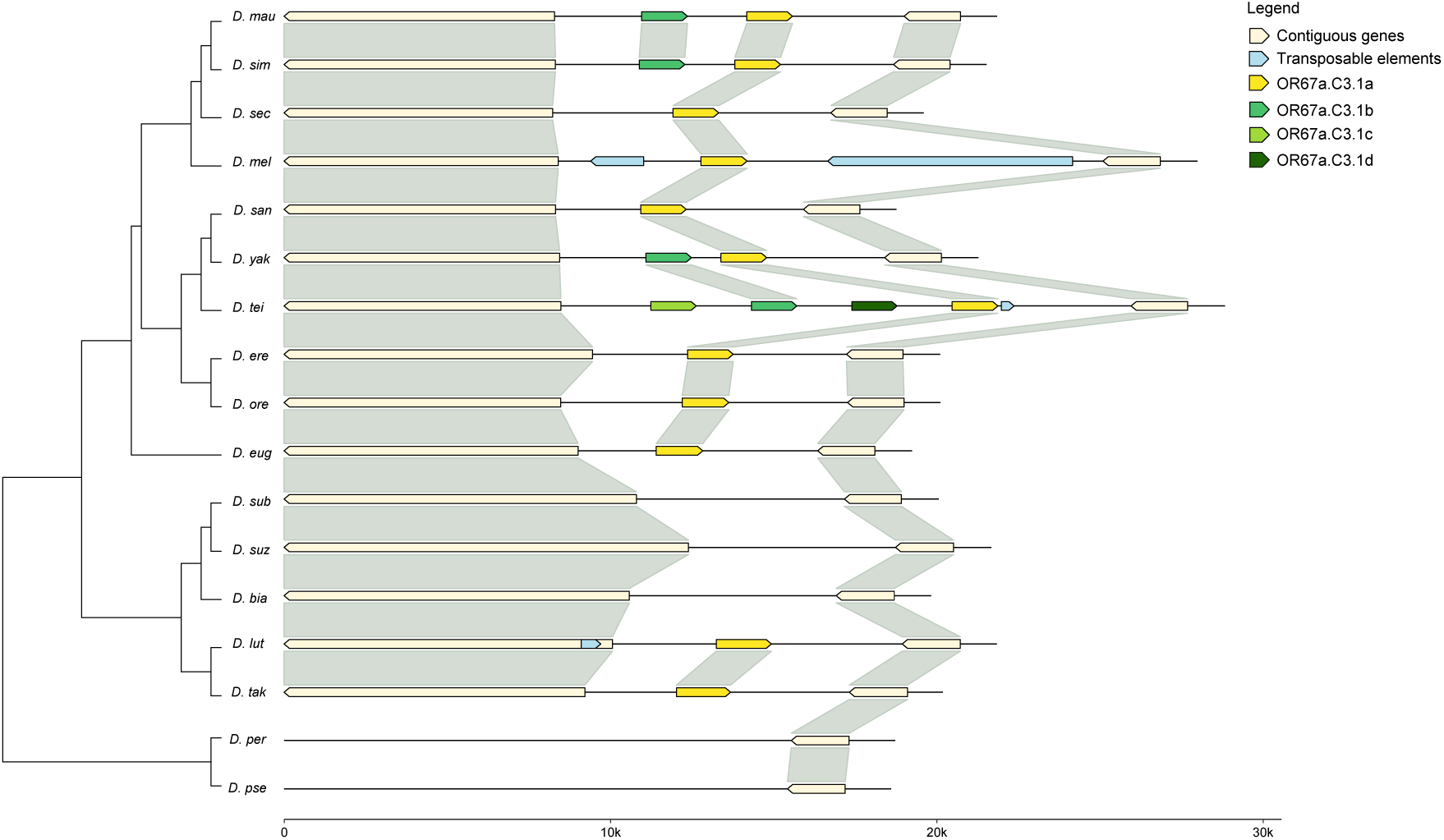
Microsynteny analysis of the *Or67.C3.1* cluster. Alignments are arranged by phylogenetic relationships (tree in left margin, not to scale). Species names are abbreviated using the first four letters of the genus and species.

**Figure S5:**
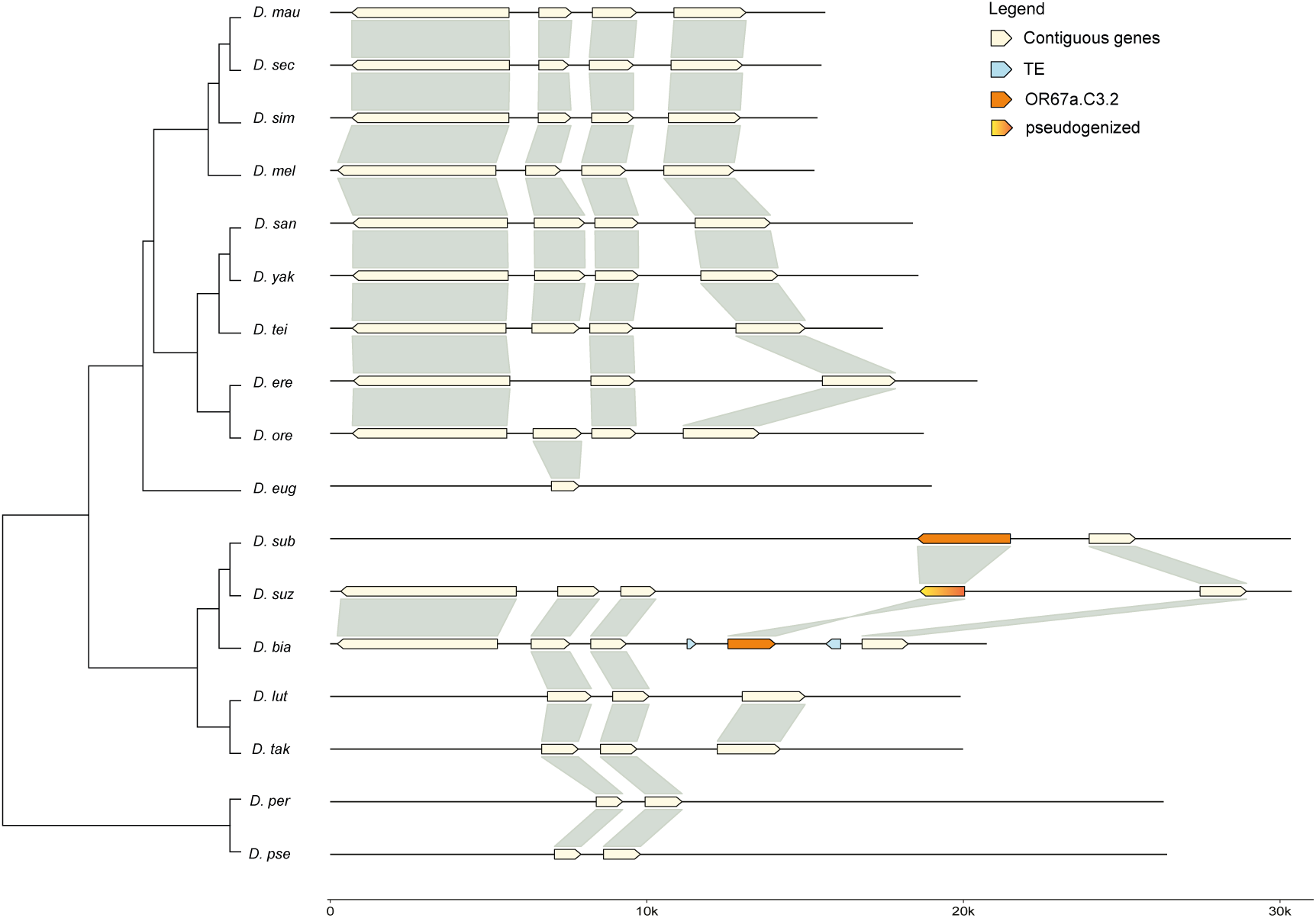
Microsynteny analysis of the *Or67.C3.2* cluster. Alignments are arranged by phylogenetic relationships (tree in left margin, not to scale). Species names are abbreviated using the first four letters of the genus and species.

**Figure S6:**
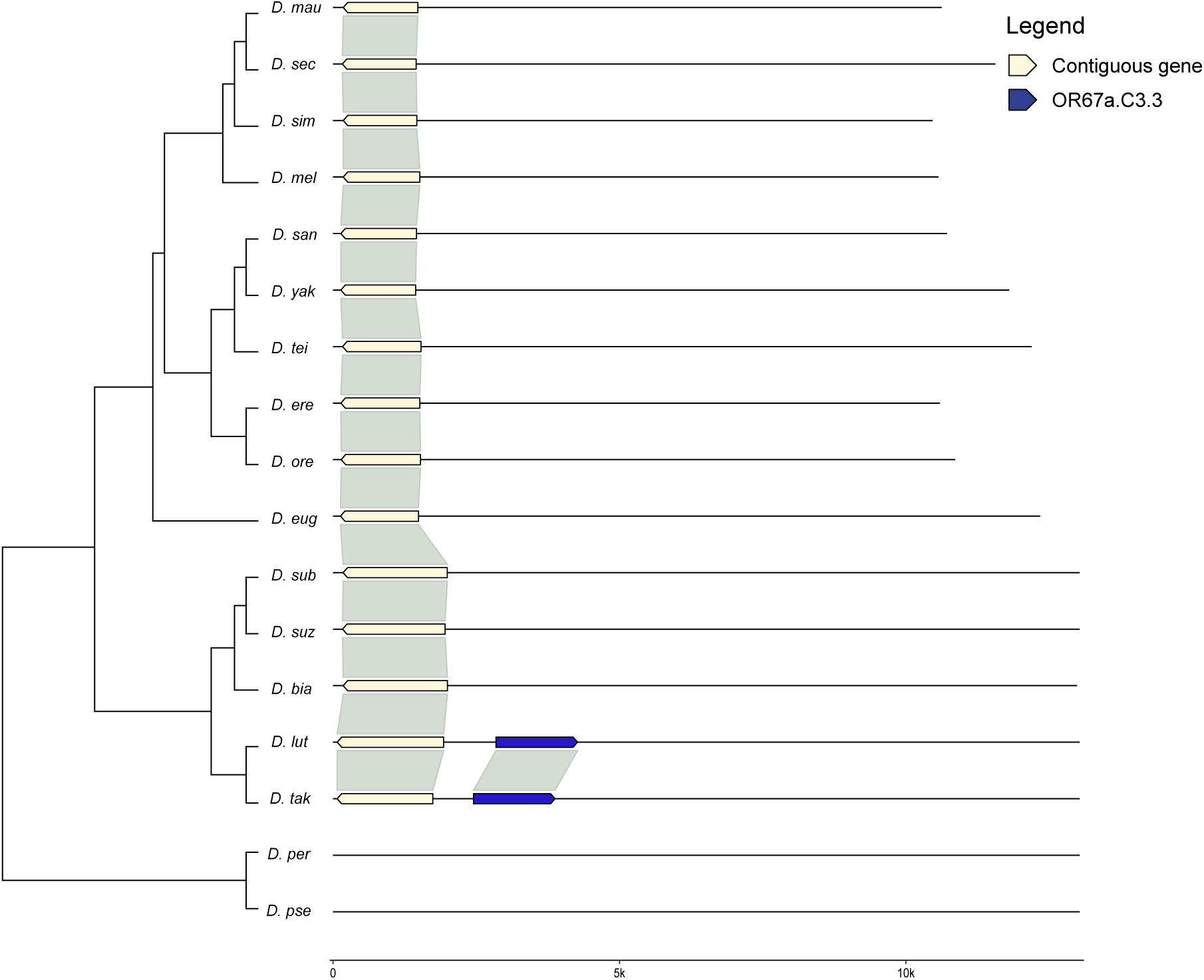
Microsynteny analysis of the *Or67.C3.3* cluster. Alignments are arranged by phylogenetic relationships (tree in left margin, not to scale). Species names are abbreviated using the first four letters of the genus and species.

**Figure S7:**
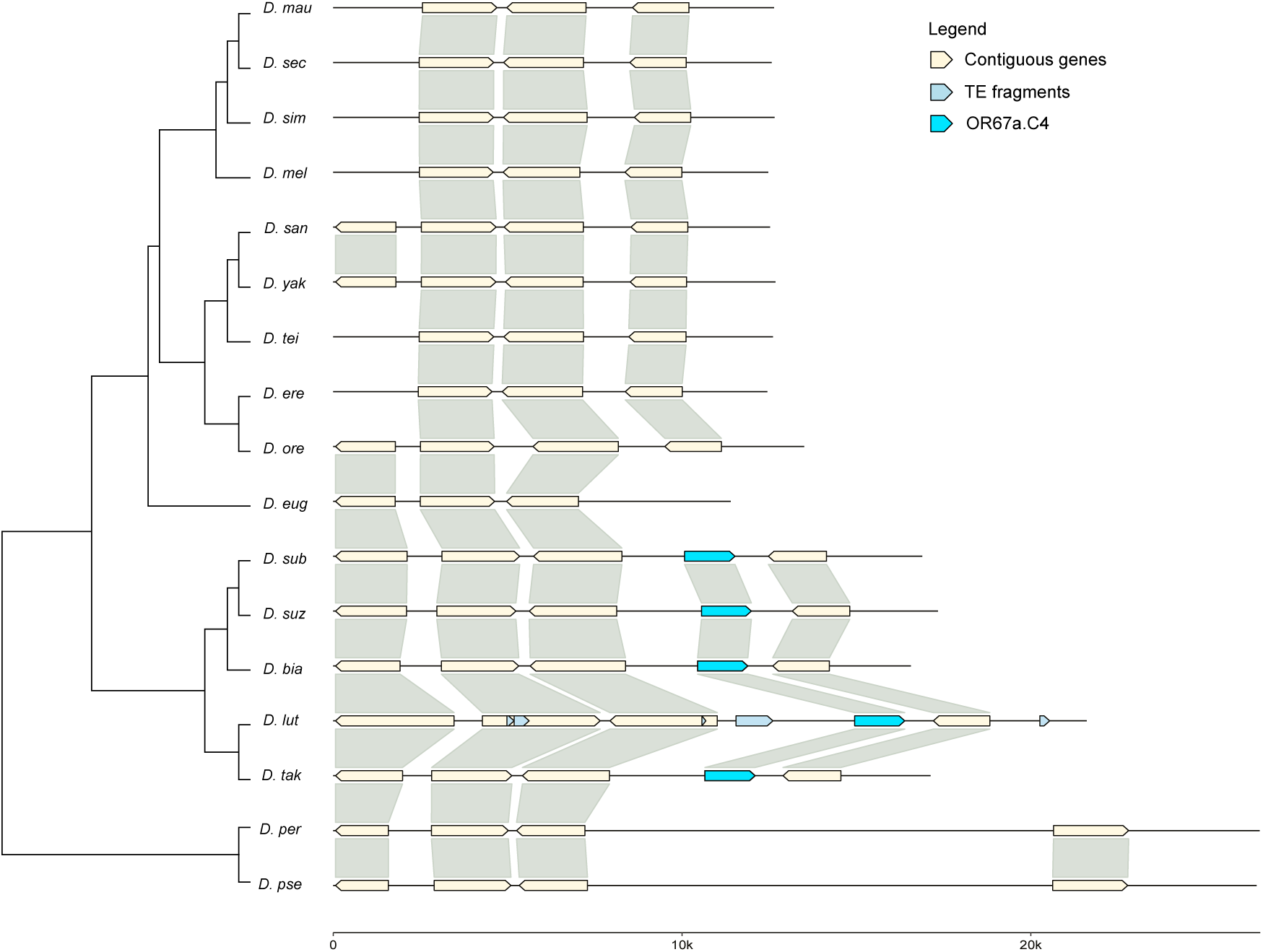
Microsynteny analysis of the *Or67.C4* cluster. Alignments are arranged by phylogenetic relationships (tree in left margin, not to scale). Species names are abbreviated using the first four letters of the genus and species.

**Figure S8:**
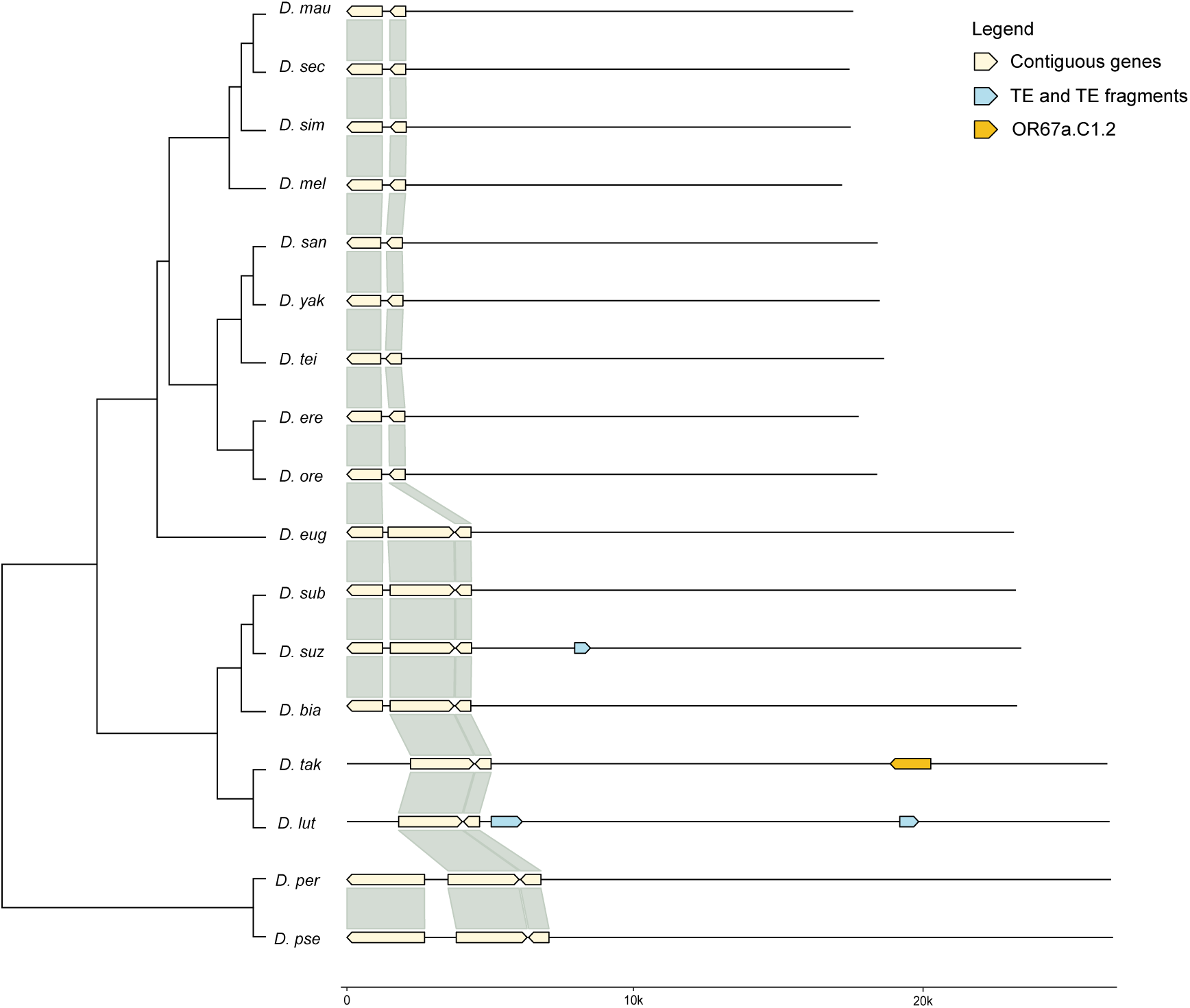
Microsynteny analysis of *Or67.C1.2*. Alignments are arranged by phylogenetic relationships (tree in left margin, not to scale). Species names are abbreviated using the first four letters of the genus and species.

**Figure S9:**
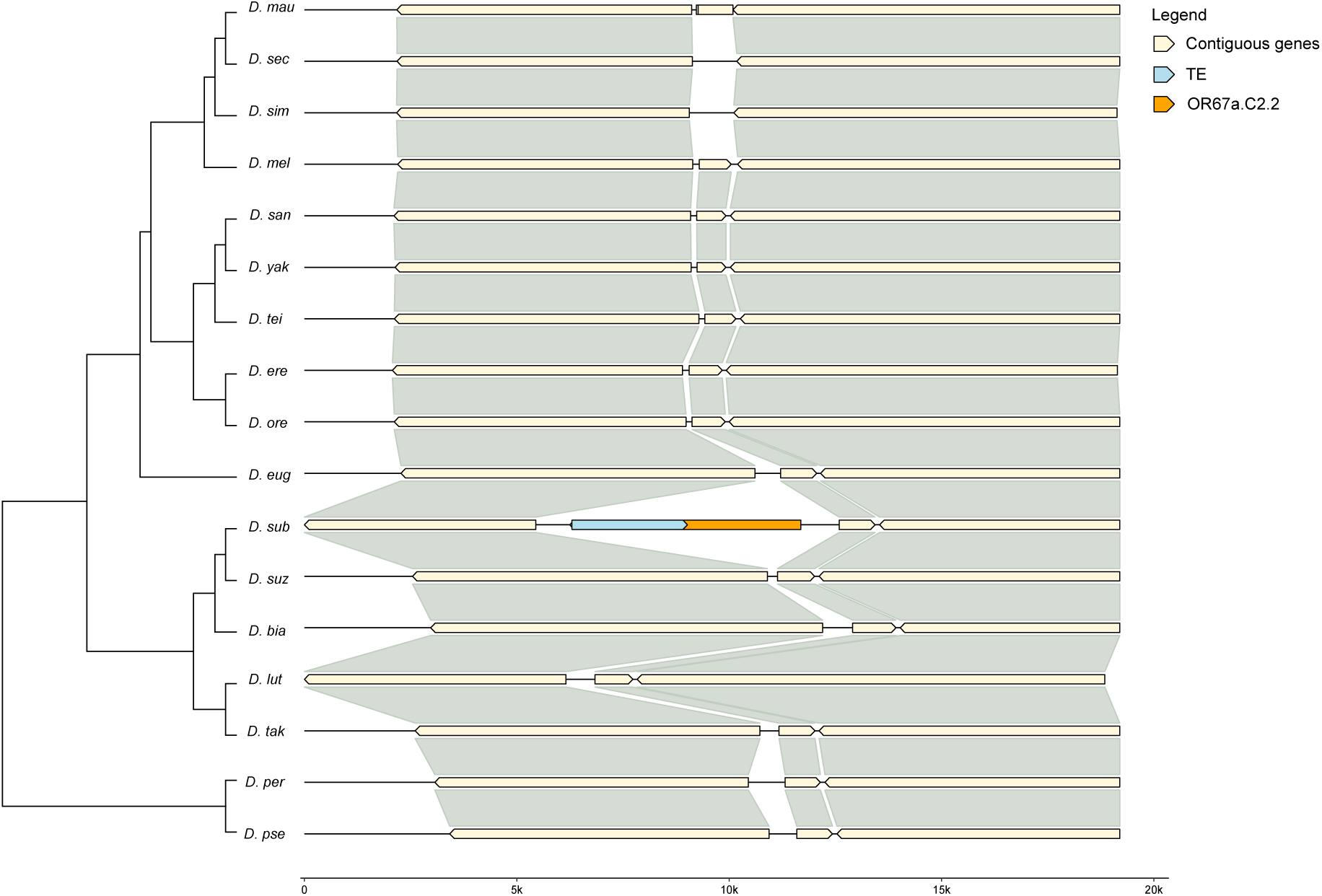
Microsynteny analysis of *Or67.C2.2*. Alignments are arranged by phylogenetic relationships (tree in left margin, not to scale). Species names are abbreviated using the first four letters of the genus and species.

**Figure S10:**
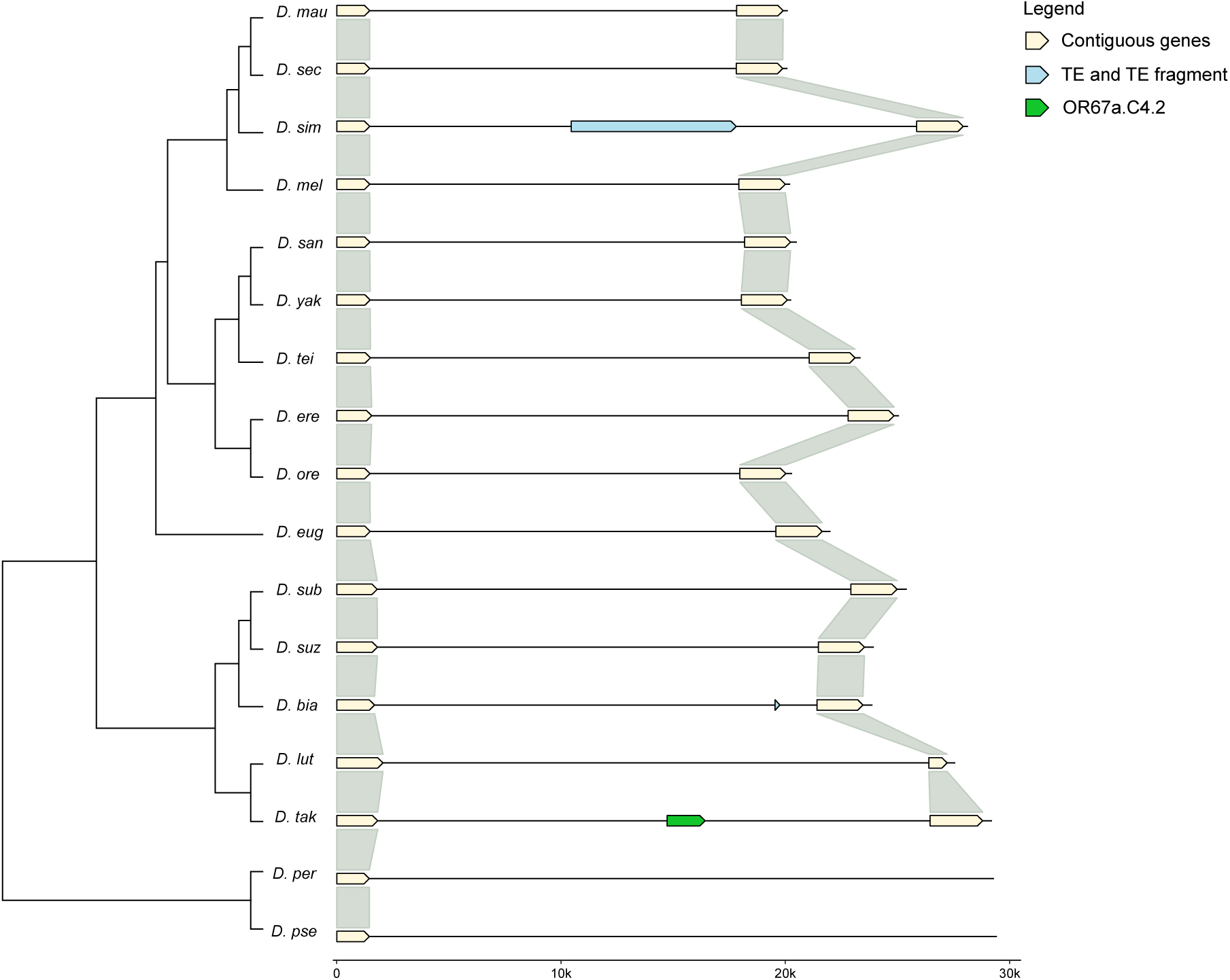
Microsynteny analysis of *Or67.C4.2*. Alignments are arranged by phylogenetic relationships (tree in left margin, not to scale). Species names are abbreviated using the first four letters of the genus and species.

**Figure S11:**
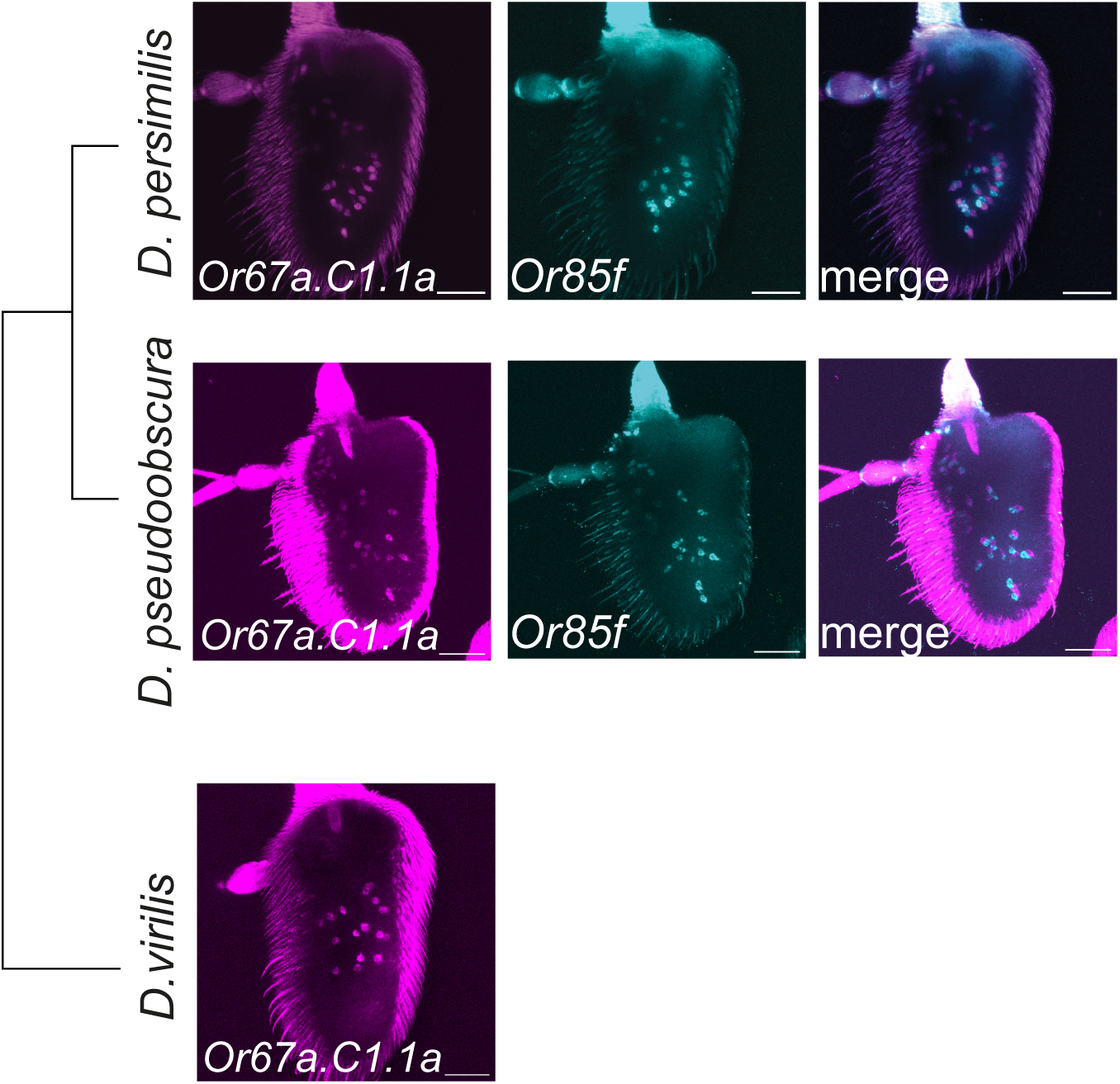
Results of co-labelling experiments using *Or67a.C1.1a* and *Or85f* HCR-FISH probes for the additional outgroup species, *D. pseudoobscura* and *D. virilis* (*D. virilis* does not have an *Or85f* ortholog). The results provide additional support to Figures 2 and 3 that the ancestral ab10A OSN population is ancestral to the species used in this study.

**Figure S12:**
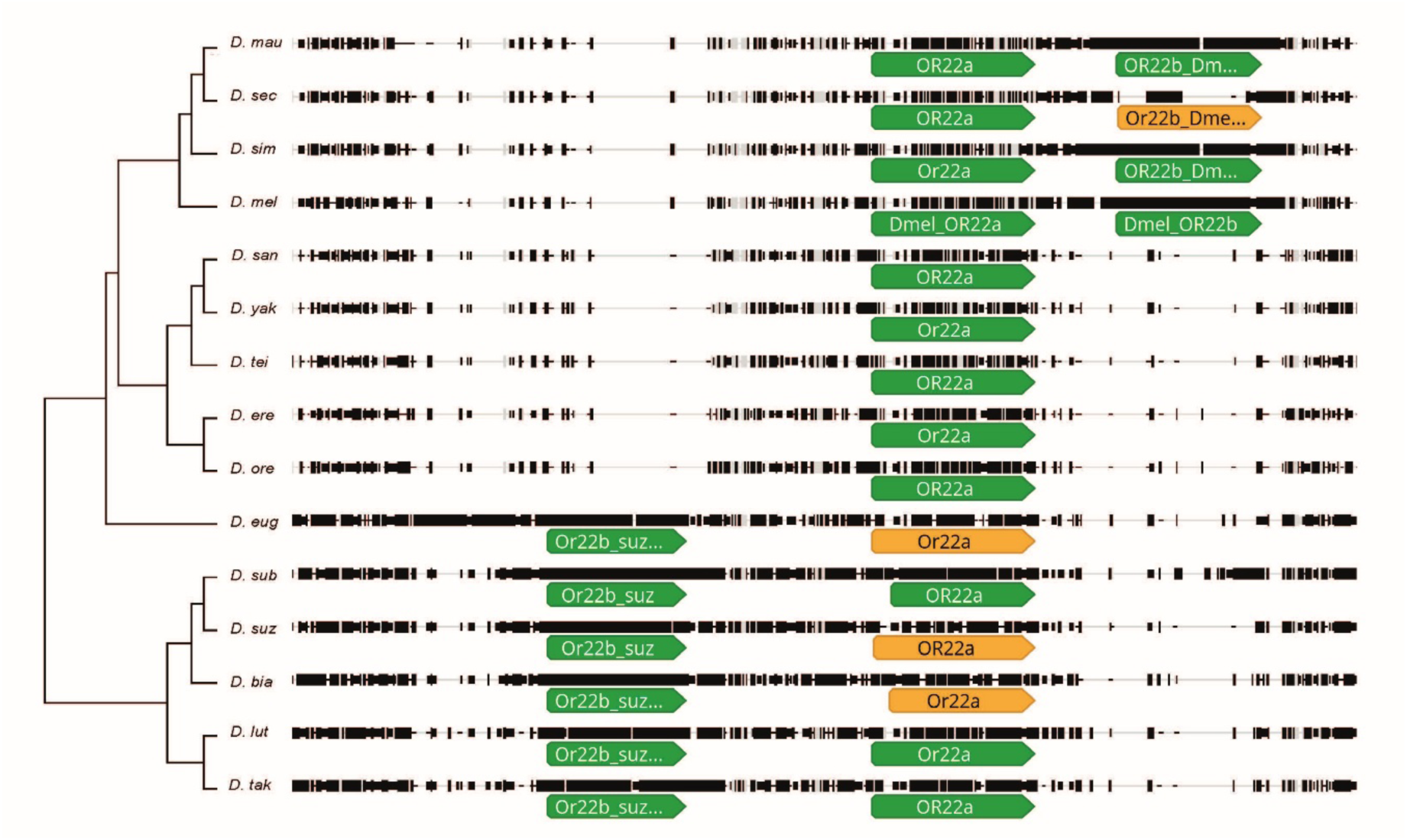
Genomic alignment of the *Or22a*-containing locus across *Drosophila* species. Species are arranged by phylogenetic relationship (tree in left margin, not to scale) and abbreviated to the first four letters of genus and species. Intact (green) and pseudogenised (orange) *Or22a* orthologs are recognisable across all species. *Or22b*-like paralogs are not shared between the two clades: *D. melanogaster* group species carry a paralog 3’ of *Or22a*, while Oriental Clade species carry a distinct paralog 5’ of *Or22a*, indicating independent duplication events in each lineage

**Figure S13:**
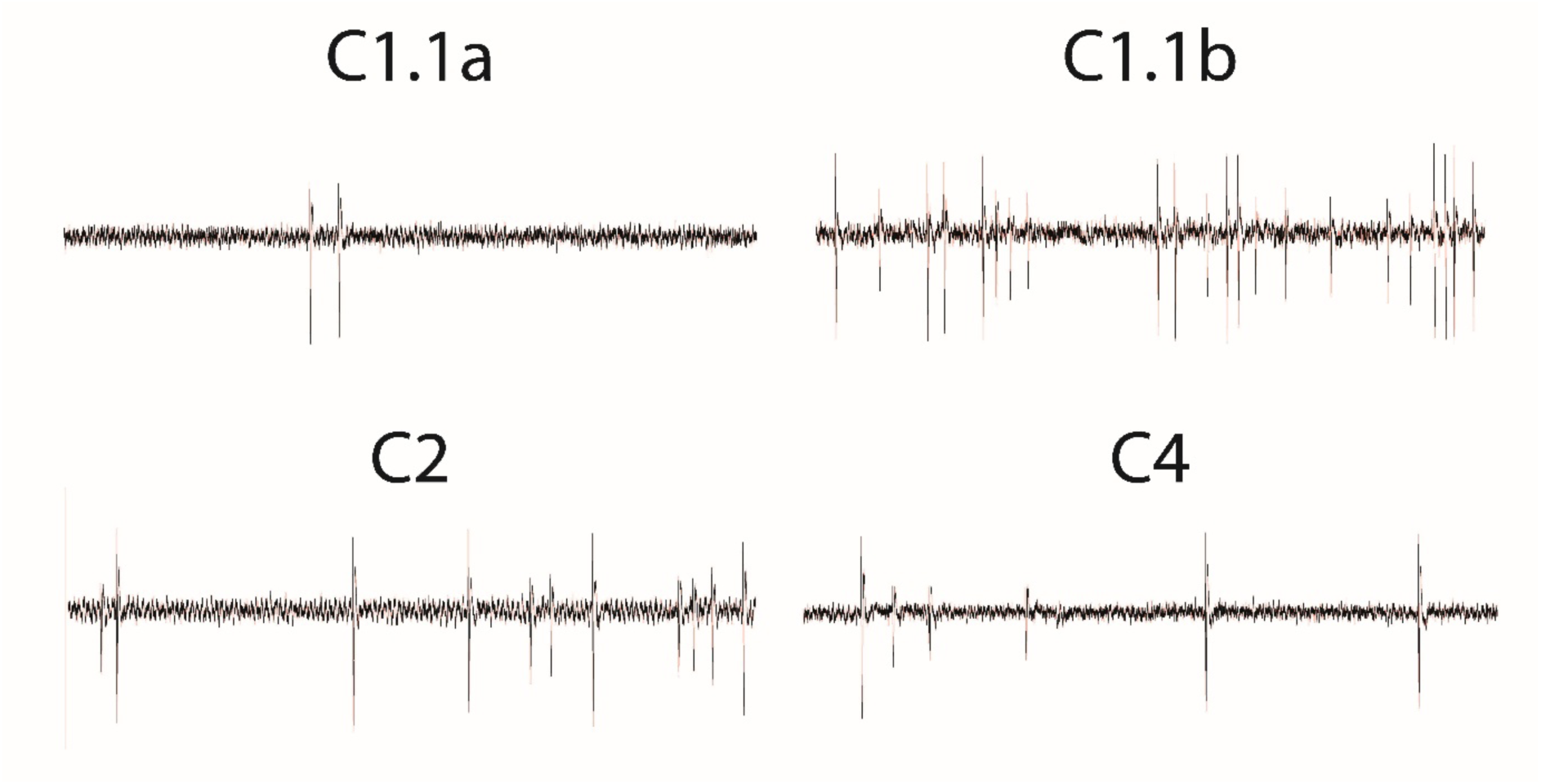
Representative single-sensillum recordings of *D. suzukii Or67a* paralogs expressed in the empty neuron system. Example spike trains illustrating spontaneous neuronal activity recorded from *D. melanogaster* ab3A olfactory sensory neurons expressing each *D. suzukii Or67a* paralog (*C1.1a*, *C1.1b*, *C2* and *C4*) prior to odorant stimulation. Neurons lack the endogenous *Or22a* receptor (*Δhalo/Cyo; Or22a-Gal4*).

**Figure S14:**
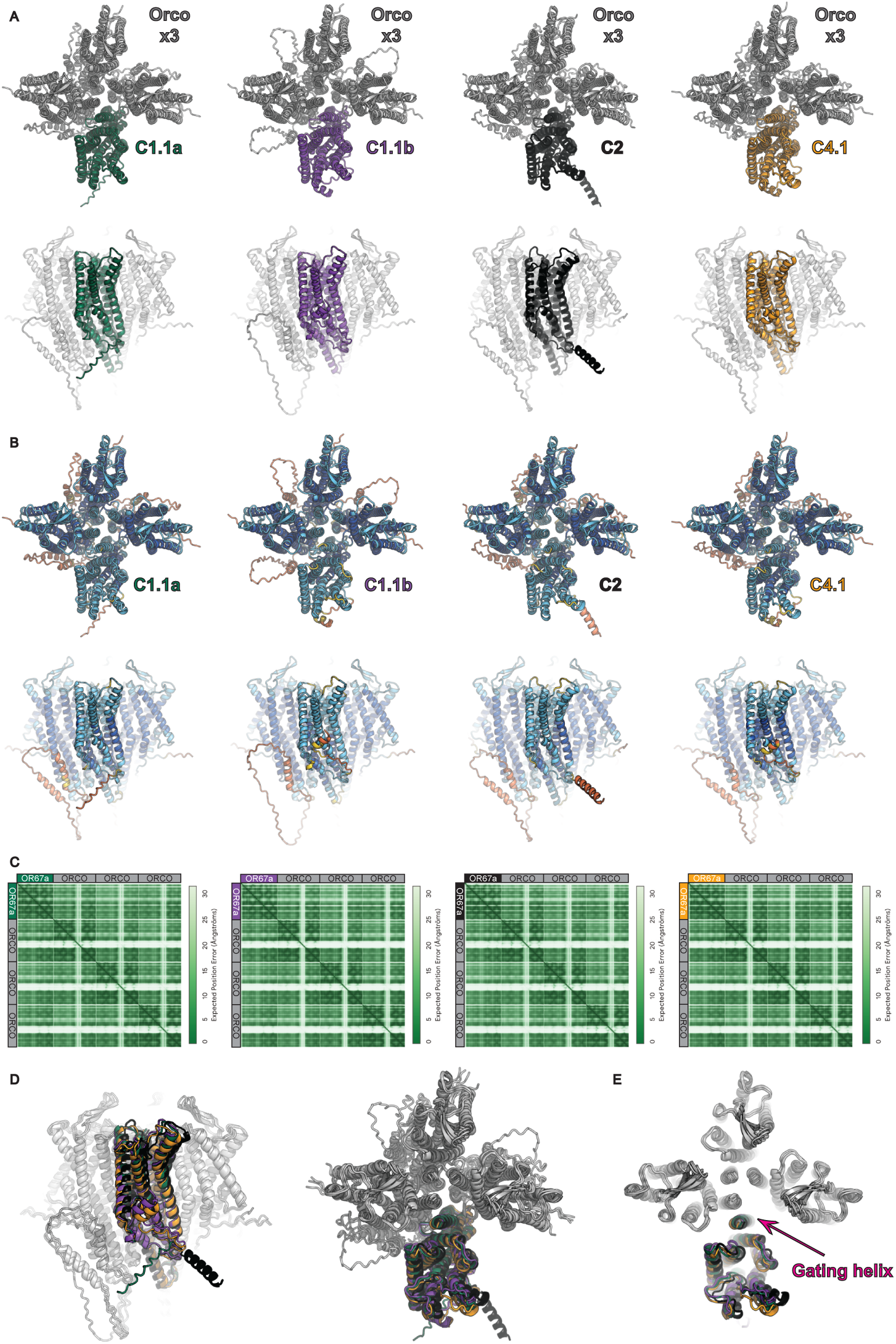
The OR67a Orco AlphaFold3 models are all predicted to form robust complexes. **A)** Orthogonal views of AlphaFold3 predicted OR67a–Orco complexes. Models are coloured by chain, there are 3 copies of a grey Orco chain per model with a single copy of OR67a C1.1a (green), C1.1b (purple), C2 (black), or C4.1 (orange). **B)** The same models coloured by pLDDT confidence: dark blue, very high (pLDDT > 90); blue, confident (90–70); yellow, low (70–50); orange, very low (< 50). **C)** Predicted Aligned Error (PAE) plots for each model, showing inter-domain and inter-subunit uncertainty. All models form a robust tetramer. **D)** Structural overlay of all four receptor–Orco complexes, demonstrating high overall similarity across predictions. Colouring as in panel A. **E)** Top-down sectional view of the overlaid models, with the gating helix highlighted, illustrating that all complexes are predicted in an open conformation.

## References

1. Benton, R. Drosophila olfaction: past, present and future. Proc Biol Sci 289, 20222054 (2022).

2. Hansson, B. S. & Stensmyr, M. C. Evolution of insect olfaction. Neuron 72, 698–711 (2011).

3. McBride, C. S. Genes and Odors Underlying the Recent Evolution of Mosquito Preference for Humans. Current Biology 26, R41–R46 (2016).

4. Carey, A. F. & Carlson, J. R. Insect olfaction from model systems to disease control. Proceedings of the National Academy of Sciences 108, 12987–12995 (2011).

5. Benton, R., et al. An integrated anatomical, functional and evolutionary view of the Drosophila olfactory system. EMBO reports 1–22 (2025) doi:10.1038/s44319-025-00476-8.

6. Herre, M., et al. Non-canonical odor coding in the mosquito. Cell 185, 3104–3123.e28 (2022).

7. Auer, T. O., Álvarez-Ocaña, R., Cruchet, S., Benton, R. & Arguello, J. R. Copy number changes in co-expressed odorant receptor genes enable selection for sensory diaerences in drosophilid species. Nat Ecol Evol 6, 1343–1353 (2022).

8. Adavi, E. D. et al. Olfactory receptor coexpression and co-option in the dengue mosquito. Preprint at 10.1101/2024.08.21.608847 (2024).

9. Auer, T. O., et al. Olfactory receptor and circuit evolution promote host specialization. Nature 579, 402–408 (2020).

10. Jové, V., et al. Sensory Discrimination of Blood and Floral Nectar by Aedes aegypti Mosquitoes. Neuron 108, 1163–1180.e12 (2020).

11. McBride, C. S., et al. Evolution of mosquito preference for humans linked to an odorant receptor. Nature 515, 222–227 (2014).

12. McBride, C. S. & Arguello, J. R. Five Drosophila Genomes Reveal Nonneutral Evolution and the Signature of Host Specialization in the Chemoreceptor Superfamily. Genetics 177, 1395–1416 (2007).

13. McKenzie, S. K. & Kronauer, D. J. C. The genomic architecture and molecular evolution of ant odorant receptors. Genome Res 28, 1757–1765 (2018).

14. Brand, P. & Ramírez, S. R. The Evolutionary Dynamics of the Odorant Receptor Gene Family in Corbiculate Bees. Genome Biology and Evolution 9, 2023–2036 (2017).

15. Mitchell, R. F., Schneider, T. M., Schwartz, A. M., Andersson, M. N. & McKenna, D. D. The diversity and evolution of odorant receptors in beetles (Coleoptera). Insect Molecular Biology 29, 77–91 (2020).

16. Matsunaga, T., et al. Evolution of Olfactory Receptors Tuned to Mustard Oils in Herbivorous Drosophilidae. Mol Biol Evol 39, msab362 (2022).

17. Legan, A. W., Jernigan, C. M., Miller, S. E., Fuchs, M. F. & Sheehan, M. J. Expansion and Accelerated Evolution of 9-Exon Odorant Receptors in Polistes Paper Wasps. Mol Biol Evol 38, 3832–3846 (2021).

18. Prieto-Godino, L. L., et al. Evolution of Acid-Sensing Olfactory Circuits in Drosophilids. Neuron 93, 661–676.e6 (2017).

19. McBride, C. S. & Arguello, J. R. Five Drosophila genomes reveal nonneutral evolution and the signature of host specialization in the chemoreceptor superfamily. Genetics 177, 1395–416 (2007).

20. Ramasamy, S., et al. The Evolution of Olfactory Gene Families in Drosophila and the Genomic Basis of chemical-Ecological Adaptation in Drosophila suzukii. Genome Biology and Evolution 8, 2297–2311 (2016).

21. Revadi, S., et al. Olfactory responses of Drosophila suzukii females to host plant volatiles. Physiological Entomology 40, 54–64 (2015).

22. Arguello, J., Fana, C., Wang, W. & Longa, M. Origination of Chimeric Genes through DNA-Level Recombination. Gene and Protein Evolution (2007).

23. Barro-Trastoy, D. & Köhler, C. Helitrons: genomic parasites that generate developmental novelties. Trends in Genetics 40, 437–448 (2024).

24. Morgante, M., et al. Gene duplication and exon shualing by helitron-like transposons generate intraspecies diversity in maize. Nature Genetics 37, 997–1002 (2005).

25. Ma, H., Wang, M., Zhang, Y. E. & Tan, S. The power of “controllers”: Transposon-mediated duplicated genes evolve towards neofunctionalization. Journal of Genetics and Genomics 50, 462–472 (2023).

26. Feuk, L., Carson, A. R. & Scherer, S. W. Structural variation in the human genome. Nat Rev Genet 7, 85–97 (2006).

27. Kapitonov, V. V. & Jurka, J. Helitrons on a roll: eukaryotic rolling-circle transposons. Trends in Genetics 23, 521–529 (2007).

28. Turissini, D. A. & Matute, D. R. Fine scale mapping of genomic introgressions within the Drosophila yakuba clade. PLoS Genet 13, e1006971 (2017).

29. Suvorov, A., et al. Widespread introgression across a phylogeny of 155 Drosophila genomes. Current Biology 32, 111–123.e5 (2022).

30. Couto, A., Alenius, M. & Dickson, B. J. Molecular, anatomical, and functional organization of the Drosophila olfactory system. Curr Biol 15, 1535–47 (2005).

31. Keesey, I. W., et al. Functional olfactory evolution in Drosophila suzukii and the subgenus Sophophora. iScience 25, 104212 (2022).

32. Takagi, S., et al. Olfactory sensory neuron population expansions influence projection neuron adaptation and enhance ssodour tracking. Nat Commun 15, 7041 (2024).

33. Matsunaga, T., et al. Odorant Receptors Mediating Avoidance of Toxic Mustard Oils in *Drosophila melanogaster* Are Expanded in Herbivorous Relatives. Molecular Biology and Evolution 42, msaf164 (2025).

34. Dekker, T., Ibba, I., Siju, K. P., Stensmyr, M. C. & Hansson, B. S. Olfactory shifts parallel superspecialism for toxic fruit in Drosophila melanogaster sibling, D. sechellia. Curr Biol 16, 101–9 (2006).

35. Pfister, P., et al. Odorant Receptor Inhibition Is Fundamental to Odor Encoding. Current Biology 30, 2574–2587.e6 (2020).

36. Hughes, G. M., et al. The Birth and Death of Olfactory Receptor Gene Families in Mammalian Niche Adaptation. Mol Biol Evol 35, 1390–1406 (2018).

37. Meyer, A. & Schartl, M. Gene and genome duplications in vertebrates: the one-to-four (-to-eight in fish) rule and the evolution of novel gene functions. Current Opinion in Cell Biology 11, 699–704 (1999).

38. Guo, S. & Kim, J. Molecular evolution of Drosophila odorant receptor genes. Mol Biol Evol 24, 1198–207 (2007).

39. Abramson, J., et al. Accurate structure prediction of biomolecular interactions with AlphaFold 3. Nature 630, 493–500 (2024).

40. Zhao, J., Chen, A. Q., Ryu, J. & del Mármol, J. Structural basis of odor sensing by insect heteromeric odorant receptors. Science 0, eadn6384 (2024).

41. Krivák, R. & Hoksza, D. P2Rank: machine learning based tool for rapid and accurate prediction of ligand binding sites from protein structure. J Cheminform 10, 39 (2018).

42. Dobritsa, A. A., van der Goes van Naters, W., Warr, C. G., Steinbrecht, R. A. & Carlson, J. R. Integrating the molecular and cellular basis of odor coding in the Drosophila antenna. Neuron 37, 827–41 (2003).

43. Prieto-Godino, L. L. et al. Olfactory receptor pseudo-pseudogenes. Nature 539, 93–97 (2016).

44. Xue, Q. & Dweck, H. K. M. Receptor sequence divergence, gain, loss, duplication, and neofunctionalization drive olfactory adaptation in *Drosophila suzukii*. Proc. Natl. Acad. Sci. U.S.A. 123, e2529586123 (2026).

45. Long, M., Betrán, E., Thornton, K. & Wang, W. The origin of new genes: glimpses from the young and old. Nat Rev Genet 4, 865–75 (2003).

46. Lynch, M., O’Hely, M., Walsh, B. & Force, A. The Probability of Preservation of a Newly Arisen Gene Duplicate. Genetics 159, 1789–1804 (2001).

47. Bontonou, G., et al. Evolution of chemosensory tissues and cells across ecologically diverse Drosophilids. 2023.04.14.536691 Preprint at 10.1101/2023.04.14.536691 (2023).

48. Mazzi, D., Bravin, E., Meraner, M., Finger, R. & Kuske, S. Economic Impact of the Introduction and Establishment of Drosophila suzukii on Sweet Cherry Production in Switzerland. Insects 8, 18 (2017).

49. Bolda, M. P., Goodhue, R. E. & Zalom, F. G. Spotted Wing Drosophila: Potential Economic Impact of a Newly Established Pest. Giannini Foundation of Agricultural Economics 13, (2010).

50. Cruz-Esteban, S. Advances in the management of Drosophila suzukii population: from olfactory and visual stimuli to development of push–pull systems. Front. Ecol. Evol. 14, 1746696 (2026).

51. Keesey, I. W., Knaden, M. & Hansson, B. S. Olfactory specialization in Drosophila suzukii supports an ecological shift in host preference from rotten to fresh fruit. J Chem Ecol 41, 121–8 (2015).

52. Li, H. Minimap2: pairwise alignment for nucleotide sequences. Bioinformatics 34, 3094–3100 (2018).

53. Sievers, F., et al. Fast, scalable generation of high-quality protein multiple sequence alignments using Clustal Omega. Molecular Systems Biology 7, 539 (2011).

54. Ronquist, F. & Huelsenbeck, J. P. MrBayes 3: Bayesian phylogenetic inference under mixed models. Bioinformatics 19, 1572–1574 (2003).

55. Chakraborty, M., et al. Evolution of genome structure in the *Drosophila simulans* species complex. Genome Res. 31, 380–396 (2021).

56. Tamura, K. Temporal Patterns of Fruit Fly (Drosophila) Evolution Revealed by Mutation Clocks. Molecular Biology and Evolution 21, 36–44 (2003).

57. Ometto, L., et al. Linking genomics and ecology to investigate the complex evolution of an invasive Drosophila pest. Genome Biol Evol 5, 745–57 (2013).

58. Katoh, K. & Standley, D. M. MAFFT Multiple Sequence Alignment Software Version 7: Improvements in Performance and Usability. Molecular Biology and Evolution 30, 772–780 (2013).

59. Camacho, C. et al. BLAST+: architecture and applications. BMC Bioinformatics 10, 421 (2009).

60. Jurka, J., et al. Repbase Update, a database of eukaryotic repetitive elements. Cytogenet Genome Res 110, 462–467 (2005).

61. Hackl, T., Ankenbrand, M., van Adrichem, B., Wilkins, D. & Haslinger, K. gggenomes: eaective and versatile visualizations for comparative genomics. Preprint at 10.48550/ARXIV.2411.13556 (2024).

62. Quinlan, A. R. & Hall, I. M. BEDTools: a flexible suite of utilities for comparing genomic features. Bioinformatics 26, 841–842 (2010).

63. Yang, Z. PAML 4: Phylogenetic Analysis by Maximum Likelihood. Molecular Biology and Evolution 24, 1586–1591 (2007).

64. Xu, B. & Yang, Z. PAMLX: A Graphical User Interface for PAML. Molecular Biology and Evolution 30, 2723–2724 (2013).

65. Yang, Z. Likelihood ratio tests for detecting positive selection and application to primate lysozyme evolution. Molecular Biology and Evolution 15, 568–573 (1998).

66. Bontonou, G., et al. Evolution of chemosensory tissues and cells across ecologically diverse Drosophilids. Nat Commun 15, 1047 (2024).

67. Zeileis, A., Köll, S. & Graham, N. Various Versatile Variances: An Object-Oriented Implementation of Clustered Covariances in *R*. J. Stat. Soft. 95, (2020).

68. Schindelin, J., et al. Fiji: an open-source platform for biological-image analysis. Nat Methods 9, 676–682 (2012).

69. Bischof, J., Maeda, R. K., Hediger, M., Karch, F. & Basler, K. An optimized transgenesis system for *Drosophila* using germ-line-specific φC31 integrases. Proc. Natl. Acad. Sci. U.S.A. 104, 3312–3317 (2007).

70. Han, C., Jan, L. Y. & Jan, Y.-N. Enhancer-driven membrane markers for analysis of nonautonomous mechanisms reveal neuron–glia interactions in *Drosophila*. Proc. Natl. Acad. Sci. U.S.A. 108, 9673–9678 (2011).

71. Stern, D. L., et al. Genetic and Transgenic Reagents for *Drosophila simulans*, *D. mauritiana*, *D. yakuba*, *D. santomea*, and *D. virilis*. G3 Genes|Genomes|Genetics 7, 1339–1347 (2017).

72. Dobin, A., et al. STAR: ultrafast universal RNA-seq aligner. Bioinformatics 29, 15–21 (2012).

73. Haese-Hill, W., Crouch, K. & Otto, T. D. peaks2utr: a robust Python tool for the annotation of 3′ UTRs. Bioinformatics 39, btad112 (2023).

74. Posit team. RStudio: Integrated Development Environment for R. Posit Software (2025).

75. Young, M. D. & Behjati, S. SoupX removes ambient RNA contamination from droplet-based single-cell RNA sequencing data. GigaScience 9, giaa151 (2020).

76. Hao, Y., et al. Dictionary learning for integrative, multimodal and scalable single-cell analysis. Nat Biotechnol 1–12 (2023) doi:10.1038/s41587-023-01767-y.

77. McGinnis, C. S., Murrow, L. M. & Gartner, Z. J. DoubletFinder: Doublet Detection in Single-Cell RNA Sequencing Data Using Artificial Nearest Neighbors. Cell Systems 8, 329–337.e4 (2019).

78. Andreatta, M. & Carmona, S. J. UCell: Robust and scalable single-cell gene signature scoring. Computational and Structural Biotechnology Journal 19, 3796–3798 (2021).

79. Gonzalez, F., Witzgall, P. & Walker, W. B. Protocol for Heterologous Expression of Insect Odourant Receptors in Drosophila. Front. Ecol. Evol. 4, (2016).

80. de Bruyne, M., Foster, K. & Carlson, J. R. Odor coding in the Drosophila antenna. Neuron 30, 537–52 (2001).

81. Hallem, E. A. & Carlson, J. R. Coding of Odors by a Receptor Repertoire. Cell 125, 143–160 (2006).

82. Benton, R. & Dahanukar, A. Electrophysiological Recording from *Drosophila* Olfactory Sensilla. Cold Spring Harb Protoc 2011, pdb.prot5630 (2011).

